# Parallel Factor Analysis for multidimensional decomposition of fNIRS data

**DOI:** 10.1101/806778

**Authors:** Alejandra M. Hüsser, Laura Caron-Desrochers, Julie Tremblay, Phetsamone Vannasing, Eduardo Martínez-Montes, Anne Gallagher

**Affiliations:** Research Center of the Sainte-Justine University Hospital, Neurodevelopmental Optical Imaging Laboratory (LIONlab), 3175 Chemin de la Côte-Sainte-Catherine, Montreal QC, Canada, H3T 1C5; Université de Montréal, Department of Psychology, Pavillon Marie-Victorin, P.O. Box 6128 Centre-ville Station, 2900 Boulevard Édouard-Montpetit, Montréal QC, Canada, QC H3T 1J4; Cuban Neurosciences Center (CNEURO), 190 e/25 y 27, Cubanacan, Playa, CP 11600, Havana, Cuba

**Author notes:** Authors contributed equally to this study (Alejandra M. Hüsser, Laura Caron-Desrochers). Equal contribution as co-senior authors (Eduardo Martínez-Montes, Anne Gallagher).

**Keywords:** Near-infrared spectroscopy, multidimensional decomposition, parallel factor analysis (PARAFAC), canonical decomposition, artifact correction, language paradigm

## Abstract

**Significance:** Current techniques for data analysis in functional near-infrared spectroscopy (fNIRS), such as artifact correction, do not allow to integrate the information originating from both wavelengths, considering only temporal and spatial dimensions of the signal’s structure. Parallel factor analysis (PARAFAC) has previously been validated as a multidimensional decomposition technique in other neuroimaging fields.

**Aim:** We aimed to introduce and validate the use of PARAFAC for the analysis of fNIRS data, which is inherently multidimensional (time, space, wavelength).

**Approach:** We used data acquired in 17 healthy adults during a verbal fluency task to compare the efficacy of PARAFAC for motion artifact correction to traditional 2D decomposition techniques, i.e. target principal (tPCA) and independent component analysis (ICA). Correction performance was further evaluated under controlled conditions with simulated artifacts and hemodynamic response functions.

**Results:** PARAFAC achieved significantly higher improvement in data quality as compared to tPCA and ICA. Correction in several simulated signals further validated its use and promoted it as a robust method independent of the artifact’s characteristics.

**Conclusions:** This study describes the first implementation of PARAFAC in fNIRS and provides validation for its use to correct artifacts. PARAFAC is a promising data-driven alternative for multidimensional data analyses in fNIRS and this study paves the way for further applications.

## 1 Introduction

Functional near-infrared spectroscopy (fNIRS) is a non-invasive neuroimaging technique that uses light of at least two different wavelengths in the near-infrared spectrum in order to assess brain activity based on neurovascular coupling. The specific absorption properties of oxygenated (HbO) and deoxygenated (HbR) hemoglobin allow individual assessments of concentration changes in both HbO and HbR separately ^1^. Although the fNIRS signal is considered to be relatively tolerant to movement ^2^, quality of data may be reduced due to abrupt changes in the light intensity caused by movement artifacts ^3^. It has been shown that the dynamics of both wavelengths provide important information for artifact detection and correction ^4^. However, current techniques for movement artifact correction (e.g. wavelet filtering, decomposition, spline interpolation, etc.) typically assume that the behavior of both wavelengths is similar in time, thus do not take advantage of the structured information offered by both wavelengths ^5–7^. 2D analyses require that data with more dimensions, such as fNIRS data, undergo superficial unfolding before processing, for example, treating both wavelengths or HbO and HbR independently. Hence, some of these 2D analysis tools are forced to impose other non-physiological constraints, such as orthogonality in the case of principal component analysis (PCA) or statistical independence for independent component analysis (ICA).

Although there are several ways to approach PCA -e.g. dimensionality reduction ^8^, classification ^9^-, from the signal decomposition point of view, PCA aims at extracting the so-called principal components, i.e. those components that explain the greatest amount of variance of the signal ^10^. It has been used for artifact removal to isolate the artifact’s signature, or to extract other relevant activities in fNIRS ^5,6,10,11^. In temporal PCA, the data is decomposed into a sum of components, each one formed by the product of two vectors: one representing the temporal principal component and the other, the corresponding topography (scores for each channel). A basic problem with PCA is that the components defined by only two signatures (time and space) are not uniquely determined. Therefore, orthogonality is imposed between the corresponding temporal signatures of the different components ^6,12,13^. Orthogonality among brain signals is, however, a rather non-physiological constraint. Even with this restriction, the extracted principal components are not completely unique, given that the arbitrary rotation of axes does not change the explained variance of the data. This has led researchers to use different mathematical criteria as the basis for choosing specific rotations (e.g. Varimax, Quartimax, Promax). In fNIRS, PCA has also been applied to target time intervals (tPCA), that is only during periods where artifacts related to articulation or other head movements occurred, instead of during the entire unsegmented signal ^3,14^. This type of targeted correction resulted in better signal quality, as compared to wavelet-based filtering and spline interpolation, while also reducing the risk of altering the signal’s global integrity ^3^. Although PCA is very common and easy to use, some authors have already discussed its pitfalls and caveats as a method for artifact correction ^7,15^.

More recently, ICA has become another popular tool for data decomposition in fNIRS ^16–18^. It has the benefit of preventing rotational freedom ^19^. However, uniqueness is achieved at the cost of imposing a constraint even stronger than orthogonality, namely, statistical independence of the temporal signatures ^20,21^. Statistical independence is appropriate for identifying artifacts with spatio-temporal signatures that are very different to those of neural activity (e.g. ocular movements), but is less appropriate for artifacts that share spatio-temporal characteristics with neural signals. What is more, ICA is commonly applied to the entire signal in contrast to a target decomposition as introduced previously for tPCA ^17^. It may thus be more challenging to achieve a satisfying correction of irregular movement artifacts. In ICA, the maximal number of components (hypothesized sources) equals the number of observations, which in neuroimaging is often rather high. It could therefore become difficult to identify the artifact’s signature, which could be split into several components.

To overcome these limitations, a multidimensional (≥ 3D) approach called parallel factor analysis (PARAFAC) ^22,23^, or less frequently referred to as canonical decomposition ^24^, could be considered as an alternative for the analysis of fNIRS data. PARAFAC is a decomposition technique applicable to any dataset that can be described in more than two dimensions (e.g. time, space, frequency, participants, conditions, signal characteristics) and allows for the extraction of different signatures present in the data. It assumes multilinear relations between the different dimensions and usually does not need any other mathematical constraints to find a unique decomposition of the data. It may therefore be used in the data pre-processing steps to isolate artifacts, as well as in the actual data analyses to extract a predominant brain activation, or other relevant characteristics of the signal. Such multidimensional decomposition was initially introduced in the field of psychometrics and linguistics as a tool for multi-factorial analysis ^22,24,25^. Over time, its use was extended to neuroimaging signals, such as the analysis of event-related potentials (ERPs) assessed by electroencephalography (EEG), in which time, space and participants were considered for the signal decomposition ^26,27^. Multidimensional decomposition with PARAFAC is not limited to 3D and could potentially be applied to data with more dimensions. In the context of ERP data, PARAFAC as a five-way analysis has successfully been used to identify differences and common characteristics of inter-trial phase coherence across conditions and subjects, including the dimensions of time, channels, frequency, subjects, and conditions ^26^. PARAFAC was also applied for data analysis in continuous EEG recordings, taking into account the temporal, spatial and frequency representation of the EEG signal ^13,28^. Indeed, using a time-frequency wavelet transformation for each channel in EEG data, Miwakeichi and colleagues ^13^ revealed PARAFAC as an appropriate tool for both the detection of ocular movement artifacts and the identification of dominant brain activation patterns. In their study, some of the artifact components identified using PARAFAC were highly similar to those extracted with principal component analysis (PCA), a well-established approach for 2D decomposition. Specifically, significant overlap was shown for eigenvalues and peaks of time/frequency components, as well as their topographical representations found with PCA and PARAFAC for activities that effectively fulfill the orthogonality requirement. PARAFAC, using the dimensions of time, space and frequency, has also been applied successfully to the EEG data of individuals with epilepsy, for the purposes of artifact detection and the identification of aberrant cerebral activation ^29^. The growing popularity of PARAFAC in neuroimaging stems from the intrinsic advantage of multidimensional decomposition, as it reflects the nature of most data gathered in neuroscience. Compared to 2D methods, the application of PARAFAC does not need to impose non-physiological constraints ^28^. What is more, since decomposition techniques represent data-driven approaches, PARAFAC could also contribute to enlighten new aspects of neuroimaging data compared to model-based techniques ^30^. Although PARAFAC has been applied for data analysis on NIRS data in the food industry ^23^, it has not yet been used in fNIRS in the field of neuroscience.

The current study aimed to introduce and validate PARAFAC as a multidimensional decomposition technique to extract and correct artifacts in fNIRS data. To account for the natural complexity and variability regarding motion artifacts, we first applied the PARAFAC technique in real cognitive fNIRS data. More specifically, data were acquired from participants that performed aloud a verbal fluency task, in which the main motion artifacts are known to be task-dependent and have characteristics that might confound the estimation of the hemodynamic response function (HRF) ^7^. PARAFAC’s performance to correct motion artifacts was compared to two traditional bi-dimensional decomposition techniques: tPCA and ICA. In doing so, we investigated differences in artifact correction efficacy related to the number of dimensions used in the decomposition techniques (i.e. treating both wavelengths as independent or as a dimension). As the true HRF was unknown and therefore did not allow to directly compare the recovered task activation to a ground truth, we used statistical analysis of correction performance based on various quality measures of the three different corrected signals.

Secondly, we investigated how artifact correction with PARAFAC performed in various controlled scenarios to disentangle in which cases a multidimensional decomposition approach that does not impose orthogonality constraints could become advantageous. For that purpose, a real artifact was extracted from a data set of the task condition and added to a clean resting-state signal. Different artifact parameters (e.g. amplitude, onset of overlapping artifacts) were controlled to produce various scenarios, including variation in the level of orthogonality among the signals combined in the simulated data. Further, the use of the resting-state signal allowed us to evaluate how artifact correction would affect the reconstruction of a synthesized HRF (HRF_sim_), which has actually been named as one of the most suitable methods to validate artifact correction ^5^. Artifact correction in these scenarios was performed with PARAFAC and tPCA only, both applied specifically during the artifacted interval (i.e. target decomposition), compared to ICA that is typically applied to the entire signal, and thus being more comparable methods. Indices for the similarity between the clean signal before adding artifacts and the signal after artifact correction, the signal’s quality as well as the recovery of the HRF_sim_ after correction, were used to compare the performance of both correction approaches.

## 2 Methods

### 2.1 Sample and data acquisition

Eighteen healthy native French-speaking adults participated in this study. One participant was excluded from the analyses, as it showed continuous noise precluding the identification of individual artifacts. The final sample for the validation of using PARAFAC for artifact correction thus included 17 participants (mean age ± standard deviation = 22.8 ± 2.0 years; 9 females and 8 males). All were right-handed and presented no neurological or psychiatric disorders. Experimental procedures were approved by the local ethics committee.

fNIRS data was acquired with a multichannel Imagent Tissue Oxymeter (ISS Inc., USA) frequency-domain fNIRS device using 14 light detectors and 60 laser light emitters, each regrouping two light sources of different wavelengths (λ_1|2_ = 690|830 nm) with an average power of 10 mW. Emitters and detectors coupled at a distance of 3 to 4.5 cm allowed for the recording of 104 channels for each wavelength. Optodes were held in place perpendicularly, using a cap that was fitted on the head of participants in accordance with the 10-20 system ^31^. Optical intensity, including information regarding the average light intensity (DC), amplitude modulation (AC) and phase shift (*φ*), was measured with a sampling rate of 19.53 Hz (Boxy, ISS Inc., USA). The channel setup covered both hemispheres equally and included the regions of interest for the investigation of language functions, i.e. frontal, temporal and parietal lobes (Fig. 1A).

**Fig. 1.**
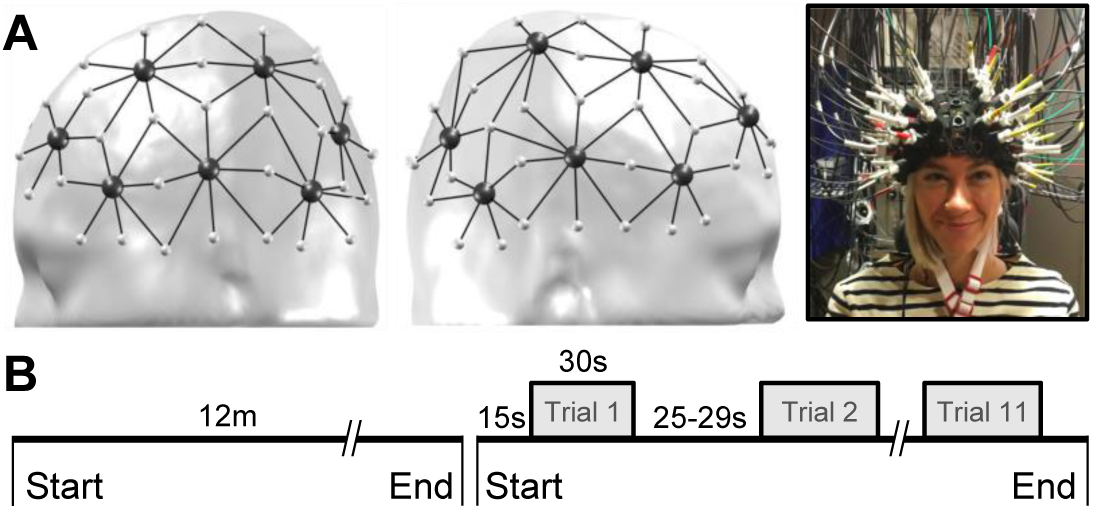
fNIRS setup: (A) Probe placement (sources in small light gray dots, detectors in dark gray dots) as shown on an adult’s head model. (B) Experimental design including a 12-minute resting-state followed by an expressive verbal fluency task that included 11 trials. Trials lasted 30 seconds and interstimulus intervals varied pseudo-randomly between 25 and 29 seconds.

Participants sat comfortably in a soundproof room. They were instructed to relax, to avoid any intentional movements or muscular tension, and to fix their gaze on the center of a screen placed at a distance of 114 cm. Participants underwent fNIRS recording during two conditions (Fig. 1B): (1) a 12-minutes resting-state with eyes open and (2) a verbal fluency task previously validated for expressive language-related activation ^32–34^. The task consisted of 11 different familiar semantic categories (e.g. animals, colors, fruits, etc.) that appeared one at a time on the screen. Participants were instructed to name as many words as possible belonging to the specified category and to continue as long as the category name appeared on the screen. We used a block design paradigm in which periods of rest (fixation cross presented on the screen) and task (semantic category) alternated (Presentations®, Neurobehavioral Systems, 2018). The interstimulus interval varied randomly between 25 and 29 seconds, while stimuli (one of the semantic categories) were always presented for 30 s. An audiovisual recording of the fNIRS session enabled visual support during offline pre-processing for the identification of movements and the timing of word articulation. Participants completed on average 10.4 ± 0.9 blocks of the verbal fluency task and named an average of 14 ± 2.2 words during each 30-s block. Data analysis was conducted with the use of a homemade toolbox (LIONirs) ^35^ adapted in SPM12 (Statistical Parametric Mapping) ^36^ in MATLAB® (The MathWorks, Inc., MA, USA).

### 2.2 Validation process

Validation of PARAFAC for artifact correction was done in two realistic applications as illustrated in Fig. 2. First, artifact correction with PARAFAC was tested on the real task-based signal of the whole sample, and its performance was compared to tPCA and ICA. Indices for the signal’s quality were used to compare the three techniques (see Sec. 2.6). Second, a simulation analysis was conducted to investigate the efficacy of PARAFAC to correct artifacts with controlled parameters. Similarity metrics, quality measures and reconstruction of the HRF_sim_ were used to evaluate artifact correction with PARAFAC and tPCA both applied in a target manner (see Sec. 2.6, 2.7 & 2.8).

**Fig. 2.**
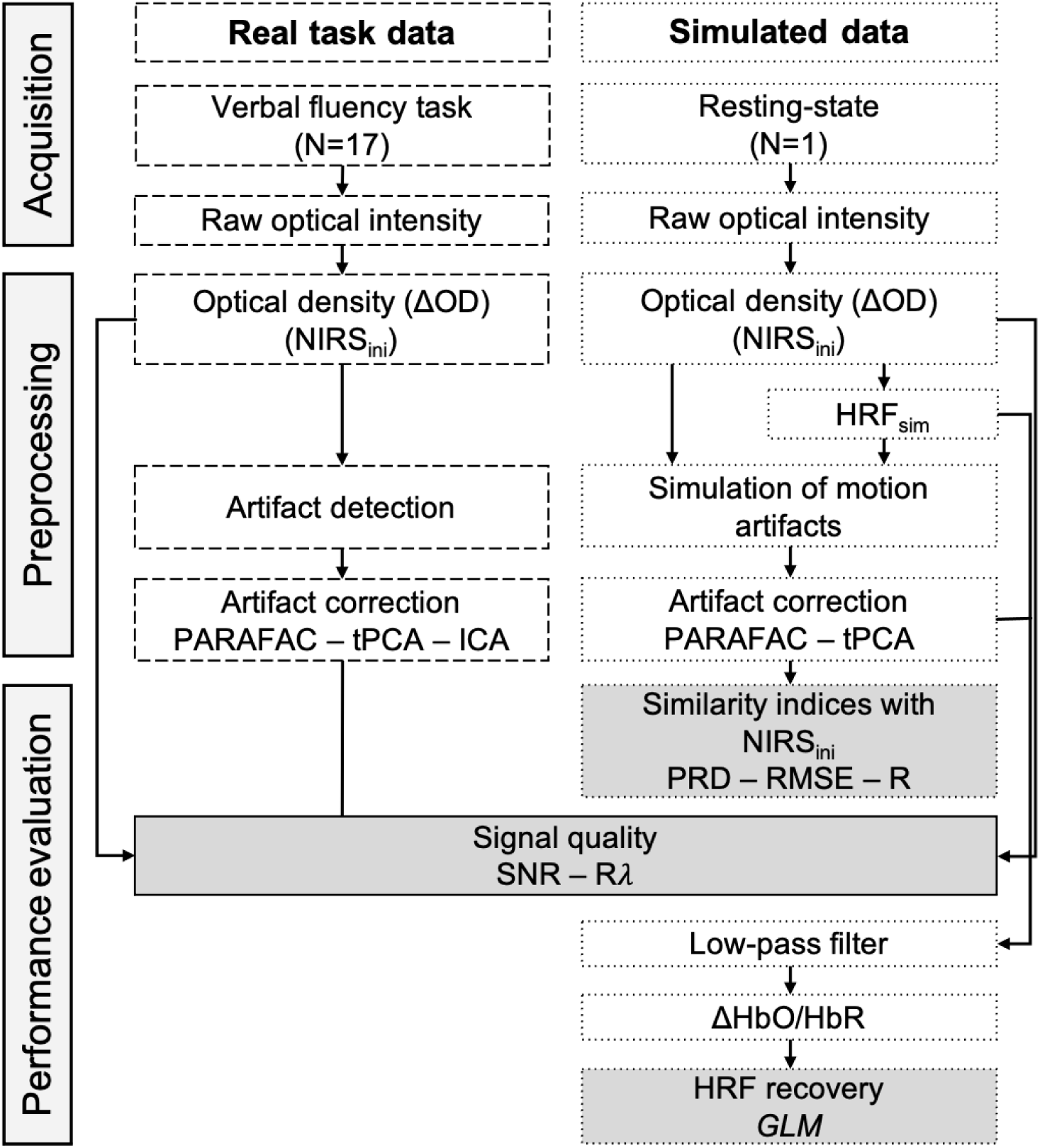
Processing streams to validate PARAFAC as a multidimensional artifact correction technique for fNIRS data. Processing of the real task data derived from 17 subjects and the simulated data based on a single-subject are indicated with dashed and dotted lines, respectively. Steps included in both are framed in a solid line. Steps in grey represent the metrics used to evaluate correction performance. Note. NIRS_ini_ = initial signal used for comparison (with artifacts for the real task data; before artifact simulation for the resting-state data). PRD = Percent root difference. RMSE = Root mean square error. R = Pearson product-moment correlation coefficient. SNR = Signal-to-noise ratio, Rλ = Pearson’s correlation between wavelengths. GLM = General linear model. HRF_sim_ = simulated hemodynamic response function.

### 2.3 Non-simulated task-related motion artifacts

Pre-processing of the task-condition data first included the automatic exclusion of channels with insufficient light intensity amplitude (average raw DC intensity across time < 100). The signal was afterwards segmented into blocks of 50 s (5 s resting-state baseline, 30 s task and 15 s resting-state), and light intensity was converted to changes in optical density (normalization of each block). We then performed a semi-automatic artifact detection using a moving-window algorithm to automatically mark segments where an abrupt change of the signal’s variance exceeded three times the average variance of the previous interval ^37^ with a window duration of 0.8 s. Events that were 2 s or less apart were considered as one and channels that were strongly correlated with a noisy channel (Pearson correlation of ≥ 0.8) within the aberrant segment were also marked as artifacted. The automatic detection step was reviewed afterwards and adjusted based on an inter-rater visual inspection of the signal’s characteristics and the video recordings. Because the task required participants to name words aloud, artifacts in the current language paradigm were mostly due to facial movements related to the muscular contraction of articulation.

### 2.4 Simulated motion artifacts

A real resting-state dataset from one of the participants in which we could identify a 180-s segment without any motion artifact was employed for the simulated artifact experiment. As for the task-based data, channels with insufficient raw light intensity amplitude were first excluded based on the previously mentioned criteria, yielding a total of 82 channels. The data was then converted to changes in optical density (normalization based on the whole 180-s segment). It served as the initial signal baseline (NIRS_ini_) for similarity and quality signal assessments.

To add a motion artifact with controlled parameters, we first used PCA to decompose all channels of a task-based fNIRS signal during a typical motion artifact (A) with a duration of 7 s.

The obtained temporal signature of the first component was then added to the NIRS_ini_. The spatial distribution of *A*, i.e. the weight of each channel was randomized, but the same distribution was added to both wavelengths with only a different overall scale. Fig. ***3*** provides an overview of the set of simulations and their parameters, while more details are provided in Fig. S1 of the Supplemental Material. For the first simulation, the amplitude of the original artifact was modulated and scaled to five different amplitudes in order to produce artifacts with various SNR (simulations 1a-1e). For the second simulation, we aimed to produce artifacts with more complex signatures (simulations 2a-f). The idea emerged from observations of artifact correction in the task-related signal and also aimed at exploring the decomposition when different artifacts are not completely orthogonal in the data. To do so, two individual artifacts (A_1_, A_2_) with varying time intervals between the onset of A_1_ and A_2_ led to different temporal orthogonality (*r*) which was evaluated for each simulation. The levels of orthogonality between the time courses of the raw signal and the first artifact *r*_*a*_ (Raw x A_1_) and between the time courses of the artifacted signal and the second artifact *r*_*b*_ ((Raw + A_1_) x A_2_), were derived from the angle between both signals (i.e. taking the time course of each signal as a vector of time points and computing the angle between both vectors). A normalized measure of the orthogonality level is then computed such that when the angle between the signals is 90 degrees, the measure is maximum and equal to 1, while when the angle departs from 90 degrees (both higher or lower) the measure linearly decreases to 0. Then, *r* values of 1 indicate perfect orthogonality between the two time courses, while *r* values lower than 1, specifically those closer to 0 indicate that the two time courses are not orthogonal to each other. Additionally, an HRF was synthesized (HRF_sim_) in SPM by the linear combination of two gamma functions as proposed in the literature ^38^. That is, gamma functions 1 and 2 had a time-to-peak of 5.4 s and 10.8 s respectively, a full-width-at-half maximum of 5.2 s and 7.36 s respectively, and a scaling coefficient for the second gamma function of 0.35. The amplitude of the HRF_sim_ was scaled by 14% for the 830 nm signal and 0.6% for the 690 nm signal, which approximately produces an HRF with an 15μM increase in HbO concentration and a 5μM decrease in HbR concentration ^5,38–40^. The response was simulated as a convolution of the HRF with a stimulus duration of 30 s resulting in a simulated physiological signal (HRF_sim_) of approximately 40 s. The HRF_sim_ was then added to the signal along with an artifact (*A*), so as to produce different overlaps between both time courses (simulations 3a-3d).

**Fig. 3.**
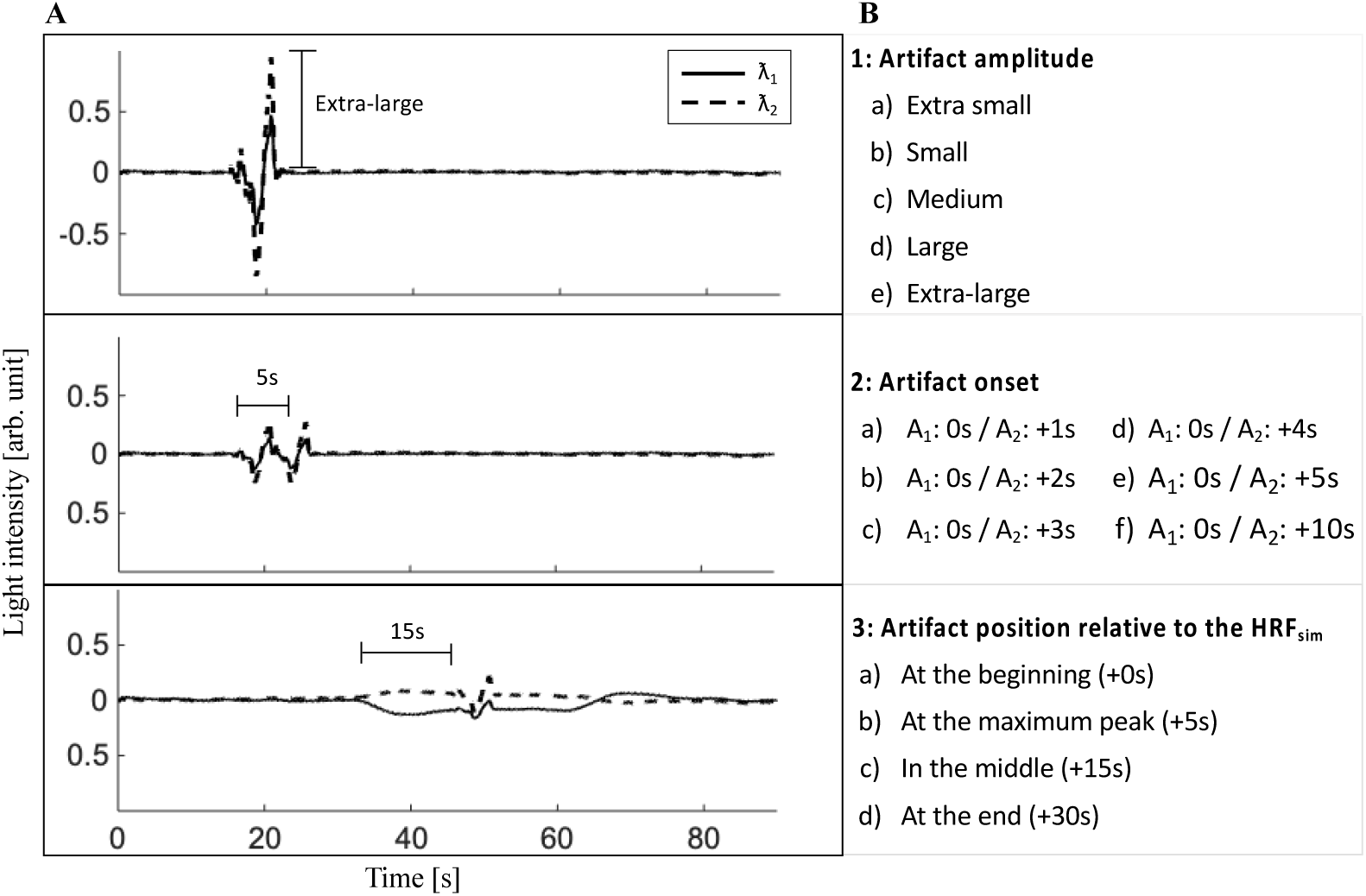
Simulated fNIRS signals: Panel A shows one example of the simulated fNIRS signal for each simulation condition 1-3. Panel B provides the details of the varying parameters of all simulations. Five scaling factors (0.4, 0.8, 1.0, 1.5, 3.0) were used to create simulated artifacts with different amplitudes. λ1|2 = 690|830 nm, A = artifact, HRF_sim_ = simulated hemodynamic response function.

### 2.5 Artifact correction

In the task-based data set, artifact correction with PARAFAC and tPCA was applied in a target manner, i.e. to the segments identified during artifact detection as described in Sec. 2.3; while, with ICA the analysis was applied to the entire continuous signal. In the simulated signals, target correction was applied to a time interval of +/-2 s around the simulated artifacts. More details on each decomposition technique will be presented in the following sections (2.5.1 & 2.5.2).

#### 2.5.1 Two-dimensional signal decomposition for artifact correction

For both ICA and tPCA, the decomposition of the 2D data matrix *X* (whose elements *x*_*ij*_ are indexed by channel *i* and time points *j*), leads to *N*_*f*_ components as defined in Eq. 1 and illustrated in Fig. 4a:

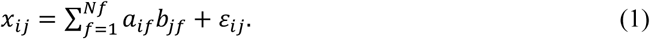

**Fig. 4.**
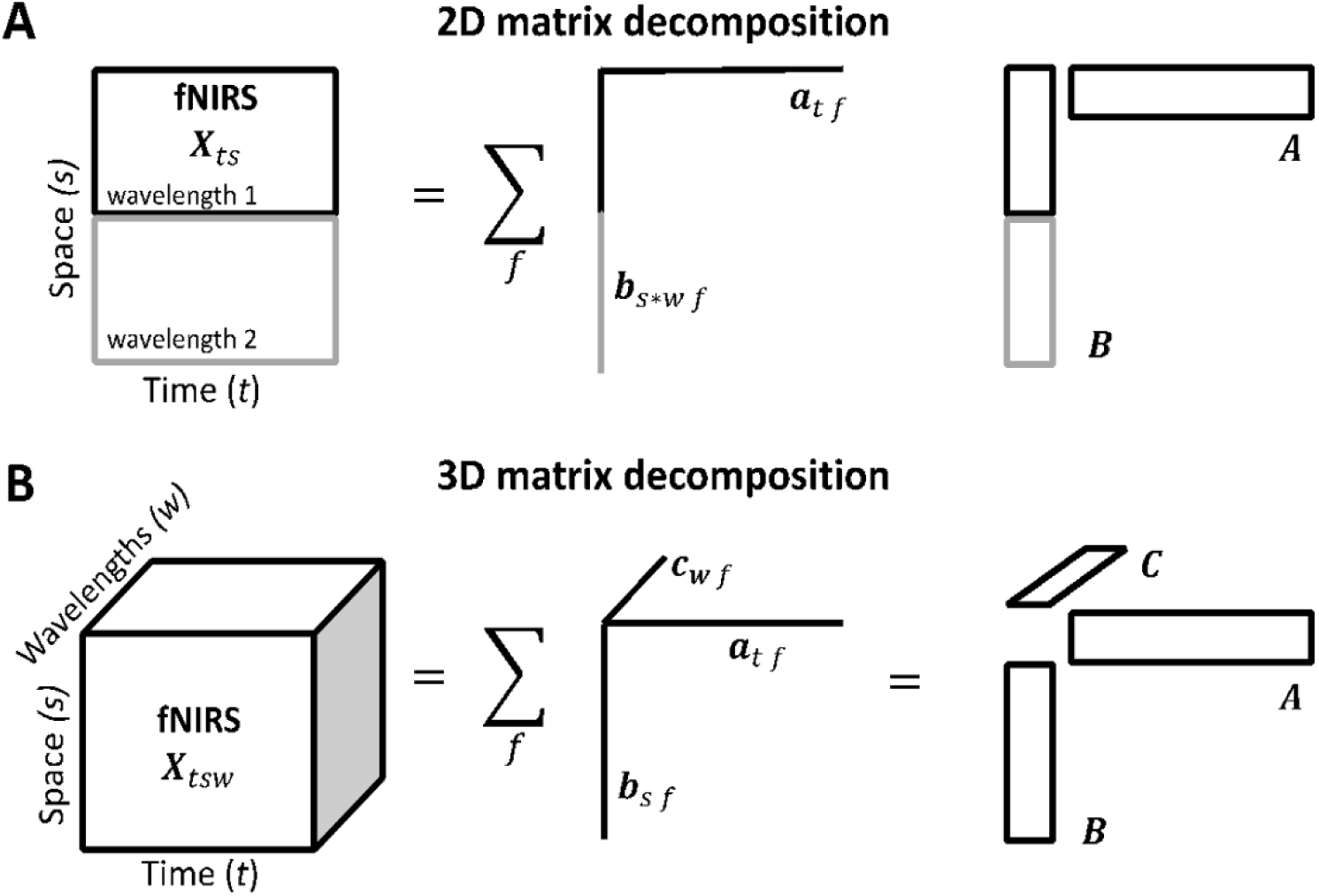
Schematic representation of the decomposition models applied to fNIRS data. **(A)** The data *X* is arranged as a 2D data array by vertically concatenating the 2D matrices with dimensions being space (s) and time (t) for each wavelength (w). tPCA/ICA decomposes the array into components, each being a bilinear product of the loading vectors representing temporal (*a*_*t f*_) and spatial signatures (*b*_*s*w f*_). The latter is formed by the spatial signatures for the different wavelengths, which are represented in components without taking into account their spatial dependence, i.e. for the same temporal signature of each component, there will be two topographies corresponding to the two wavelengths. Matrices *A =* {*a*_*f*_} and *B =* {*b*_*f*_}, contain as columns the temporal and spatial signatures for all components, respectively. **(B)** The data *X* is arranged as a 3D data array with dimensions being time (t), space (s) and wavelengths (w). PARAFAC decomposes this array into the sum of components, each being a trilinear product of loading vectors representing temporal (*a*_*t f*_), spatial (channel, *b*_*s f*_) and spectral (wavelength, *c*_*w f*_) signatures. In practice, the decomposition consists of finding the matrices *A =* {*a*_*f*_}, *B =* {*b*_*f*_} a*nd C =* {*c*_*f*_} that explain *X* with minimal residual error.

The matrix concatenates two separate data matrices (*X*_1_, *X*_*2*_) corresponding to the 2D structure for both wavelengths. The maximum number of components *N*_*f*_ could be equal to or less than the smaller dimension of the *X* matrix dimension, i.e. twice the number of channels or number of time points. Each component is modeled as the product of two factors/vectors which represent signatures of the space (*a*_*f*_) and time (*b*_*f*_) dimensions. The temporal signatures are constrained to be orthogonal among components, and rotated in order to obtain those signatures that offer the highest variance explanation (varimax) from all the infinite solutions of the decomposition. The unexplained part of the data is considered irrelevant activity or noise (*ε*).

Artifact correction with tPCA was performed on each time interval containing artifacts as specified by the time interval of the simulated artifacts or the identified artifact detection of the task-based data set. From the obtained components we subtracted the first, which explains most of the variance based on the assumption that the highest variance in the data for each segment is assumed to be caused by the artifact.

Artifact correction with ICA, was done using BrainVision Analyzer (Brain Products GmbH, Gilching, Germany), and followed the steps proposed by Plank ^41^. First, optical density fNIRS data were exported and ICA was applied on the entire unsegmented signal, as for the method commonly reported in the literature ^17^. Noisy intervals reflecting artifacts were identified by semi-automatic inter-rater artifact detection (see Sec. 2.3). We visually selected the ICA components reflecting the artifact, based first on their time course, i.e. high variation of light intensity similar to the artifact signature. Secondly, we rejected those whose subtraction would induce artificial artifacts elsewhere. Data was subsequently imported back into the LIONirs toolbox ^35^ in order to apply segmentation (see Sec. 2.3).

#### 2.5.2 Multidimensional signal decomposition

The fNIRS data naturally offers the time courses of all channels for two wavelengths, i.e. two separate data matrices. This data can be arranged in a tri-dimensional array, as illustrated in Fig. 4b. The dimensions of this 3D array are time (indexed by the time points in the analyzed segment), space (indexed by channels) and wavelength (indexed by the two wavelengths). As explained above, PARAFAC establishes a trilinear decomposition of each element of the fNIRS data array (*X*_*tsw*_) in *N*_*f*_ components, each being the product of three factors^5^ as defined in Eq. 2:

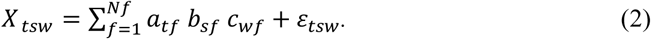

The estimated factors, *a*_*tf*_, *b*_*sf*_ and *c*_*wf*_ are the elements of the so-called loading matrices *A, B and C*, whose column vectors *a*_*f*_ *=* {*a*_*tf*_}, *b*_*f*_ *=* {*b*_*sf*_ }, *c*_*f*_ *=* {*c*_*wf*_}, represent the temporal, spatial and wavelength signatures of each component. The main advantage of this method is that it provides a unique decomposition of the fNIRS data into components reflecting different activities that do not need to be orthogonal or statistically independent in any of the dimensions. As long as an activation shows a different behavior in one of the dimensions, it can be extracted as a separate component. In this application, the spatial dependency between wavelengths is therefore exploited in order to perform the decomposition. The uniqueness of the solution is guaranteed when the number of components (*N*_*f*_) is smaller than the sum of the ranks of the three loading matrices. In the case of noisy data, it is very likely that the loading matrices are always full rank. Uniqueness is thus guaranteed as long as the number of components (*N*_*f*_) is smaller than half the sum of the number of time points (*N*_*t*_), number of channels (*N*_*c*_), and number of wavelengths 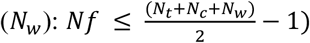. For instance, even in a small array of 20 time points and eight channels at two different wavelengths, a unique decomposition could be achieved, using up to 14 components 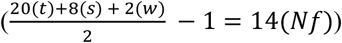. Computationally, the decomposition is achieved by the Alternating Least Squares algorithm, as it has been used in previous studies on neuroscience data ^13,28,29,42^. The only indeterminacies in the least-squares solution are trivial and easy to handle: the order of the additive components and the relative scaling of the signatures ^23^.

In this work, we used the PARAFAC implementation from the N-way Toolbox ^43^, which was included into the LIONirs toolbox ^35^ so as to visualize and apply the decomposition exclusively during selected time intervals and channels. Components were ordered according to their importance in explaining the data’s variance (similar to tPCA). The scale of the data was kept in the temporal signatures, while the other dimensions were normalized so as to have Frobenius norms equal to one. Since measures from both wavelengths are sampled simultaneously at each specific position on the scalp, movement artifacts affect their amplitudes similarly. During time intervals containing artifacts, the time courses of the signal from the two wavelengths commonly show a drastic and correlated increase as compared to the task-related or baseline signal ^4^. By using multidimensional PARAFAC decomposition, we can take advantage of this information for the adequate selection of components of the artifact’s signature. PARAFAC decomposition was specified to extract between two to four components, allowing a clear separation of the artifact signatures representing the artifacts characteristics from the rest of the signal. The appropriate components were selected based on (1) a visual inspection of their temporal overlap with the artifact, (2) the smallest number of possible components that would sufficiently correct the artifact, and (3) components showing similar weights for both wavelengths. In simulations, a standardized PARAFAC decomposition with 3 components was applied. Two components that clearly showed evidence of an amplitude change typically related to a movement artifact, i.e. short impulses, were discarded as reflecting the artifact in the original signal.

### 2.6 Signal quality

The signal’s overall quality was estimated by two quality measures ^4,5,44^:

1. <u>Signal-to-noise-ratio:</u> For each data segment with an artifact we estimated the signal-to-noise ratio (SNR), as defined by Sweeney and colleagues ^44^:

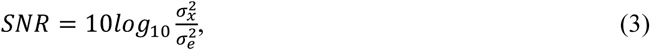

where 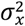 represented the signal’s variance computed in segments of data without an artifact. In case of the real task-based data set, 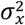 was computed in a 5-s interval during the baseline of the task condition. For the simulations, 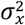 was computed in a 5-s interval at the beginning of the resting-state signal. 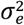 represented the variance of a segment where an artifact was identified or simulated. Whenever an artifact contaminated the signal, 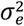 would usually be higher than 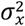 because it comprised both the variance of the physiological signal and the variance of the artifact. It was therefore expected that the SNR of the signal before artifact correction would be negative. After artifact correction, a negative SNR (σ_x_ < σ_e_,) would indicate that the artifact had not been entirely removed, a positive SNR (σ_x_ > σ_e_,) would imply overcorrection meaning that artifact correction removed the artifact as well as parts of the relevant physiological activity, and an SNR of about 0 (σ_x_ = σ_e_) would suggest that the artifact had been eliminated to the extent that the variance of the corrected signal would not differ from the baseline segment without an artifact. In case of the task-based data set, the SNR of the uncorrected signal served as a baseline, thus the more the SNR of the corrected signal differed from the baseline value and approached 0, the better the artifact had been corrected. Since the ground truth of the signal’s variance was unknown and besides the artifact also the hemodynamic response may contribute to the signal’s variance, no further interpretation of a negative or positive SNR would have been appropriate. For the simulations, the SNR of the initial signal before artifact simulation (NIRS_ini_ and NIRS_ini_ + HRF_sim_) served as a reference and the more the SNR of the corrected signal would approach this value, the better the performance of artifact correction was assumed to be.
2. <u>Pearson’s correlation coefficient (Rλ)</u> between the time courses of both wavelengths from the same site were used as a subsequent quality measure to evaluate artifact correction performance ^4^. This was based on the assumption that a high correlation coefficient indicates the presence of artifactual signals measure simultaneously by both wavelengths, since in a clean signal these temporal courses appear much less correlated. Similar to how it was previously done for the SNR, we computed Rλ for all segments with identified or simulated artifacts before (i.e. uncorrected artifact reference) and after artifact correction (i.e. corrected signal). For the task-based data, Rλ was additionally calculated for artifact-free baseline intervals (i.e. artifact-free reference) and in case of the simulations for the intervals of the initial signal (NIRS_ini_ and NIRS_ini_ + HRF_sim_) before artifact simulation (i.e. artifact-free reference).

### 2.7 Similarity indices

For the analysis of simulated data, three additional metrics were used to evaluate performance of artifact correction: (1) the percent root difference (PRD), (2) the root mean square error (RMSE) and (3) the Pearson product-moment correlation (R). These allowed us to estimate the degree of correspondence between the initial signal without simulated artifacts (NIRS_ini_ and NIRS_ini_ + HRF_sim_) and the signal after correction of the simulated artifacts ^40,45,46^. PRD, RMSE and R were defined as follows:

1. The percent root difference (PRD)

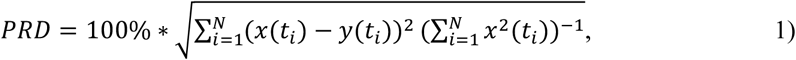
2. Root mean square error (RMSE)

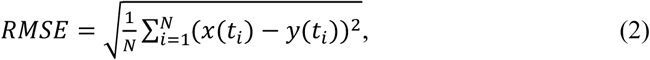
3. Pearson product-moment correlation (R)

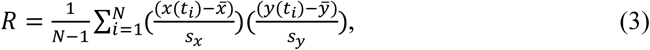

where 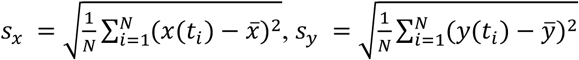.

For all three indices, *x*(*t*_*i*_) and *y*(*t*_*i*_) represented the i-th point of the time courses of the corrected signal and of the initial artifact-free signal (NIRS_ini_ and NIRS_ini_ + HRF_sim_), respectively. *N* corresponded to the duration of the time courses and 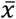 and 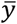 to their respective mean values along time. An ideal artifact correction would have been achieved when the artifact was completely removed, and the corrected signal maximally resembled the initial signal. As PDR and RMSE inform on the difference of the two signals, the smaller they were, the more accurate the performance of artifact correction was. On the contrary, the R index represents the similarity of both signals, hence a higher value suggested better artifact correction.

### 2.8 HRF recovery

In order to investigate how artifact correction would affect the interpretation of the hemodynamic response, in the last simulations (3a-d) we aimed to recover a previously synthesized HRF. Therefore, a Butterworth low-pass filter (filter order = 4, cut-off frequency = 0.2 Hz) was applied to the signal to remove oscillations caused by heartbeat and respiration. The optical intensity changes of the two wavelengths were then transformed into relative concentration changes of HbO and HbR, done so by using an age-adapted differential pathlength factor (DPF) and the modified Beer-Lambert Law ^47,48^. A general linear model (GLM) with the HRF_sim_ as a predictor was then applied. The GLM assumes that a linear relation exists between different inputs ^49^, and has already been applied in several fNIRS studies to specify the HRF ^6,50,51^. The cerebral activation of the signal (*y*) is defined as:

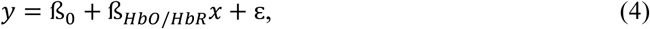

Where *y* referred to the analyzed signal, ß_0_ was a constant, ß_HbO/HbR_ represented the predictive value of the simulated hemodynamic response represented by *x* (HRF_sim_) for both concentration changes of HbO and HbR, respectively, and ε represented the error or the unexplained part of the signal. GLM was applied to a 60-s interval that was set to 15 s before and 45 s after the onset of the HRF_sim_. The GLM allowed us to estimate how much of the signal’s variance (R^2^) could be predicted by the HRF_sim_. This estimation was conducted for the initial signal without artifacts (NIRS_ini_ + HRF_sim_), the uncorrected signal with a simulated artifact, and for the corrected signal.

### 2.9 Statistical analysis

Quality and similarity measures were computed for each channel separately and then averaged across channels for statistical analysis. Prior to statistical analysis, correlation coefficients (R and Rλ) were standardized using Fisher’s transformation to obtain values following a normal distribution. For the real task-based data, we conducted statistical analysis to compare the signal quality metrics among the motion artifact correction techniques. A repeated measures ANOVA including PARAFAC, tPCA, ICA and the uncorrected reference signal as the within factor, was performed independently for the mean of each metric across all channels (i.e. SNR, Rλ). In case of the Rλ, the reference of the baseline (non-artifacted signal) was included as another condition. Follow-up paired contrasts were conducted with a critical alpha of 0.05. For the simulations, we applied a repeated measures ANOVAs to compare the mean outcome of the quality and similarity metrics between the different simulations, i.e. 1a-e for the amplitude scaling, 2a-2f for the complex artifacts and 3a-d for the HRF_sim_, and within correction condition, i.e. PARAFAC, tPCA, and the uncorrected signal. Tukey correction for multiple comparisons was applied for post-hoc analysis.

## 3 Results

### 3.1 Correction of non-simulated task-related motion artifacts

First, decomposition with PARAFAC was applied for the correction of real task-based motion artifacts. The signal’s quality (SNR, Rλ) after correction with PARAFAC was evaluated and compared to the signal before correction (Uncorrected), the signal after correction with two 2D decomposition techniques (i.e. tPCA and ICA), and a reference signal of a segment without artifacts (Rλ only).

Fig. **5** provides an example of target artifact correction with PARAFAC illustrating the 3D decomposition for a motion artifact detected in the real task-based signal. The time course of the three components obtained with PARAFAC allowed to differentiate between the two components with distinct artifact signatures, and the one component representing the clean signal. Spatial distribution of each component is shown as a loading matrix and helps to identify in which channels (localization) the artifact’s signatures were most important. The two components that appear to represent the artifact showed similar scores for both wavelengths, as represented in the wavelength signatures. Subsequently, removal of PARAFAC first and second components resulted in a less noisy time interval and satisfyingly corrected signal.

**Fig. 5.**
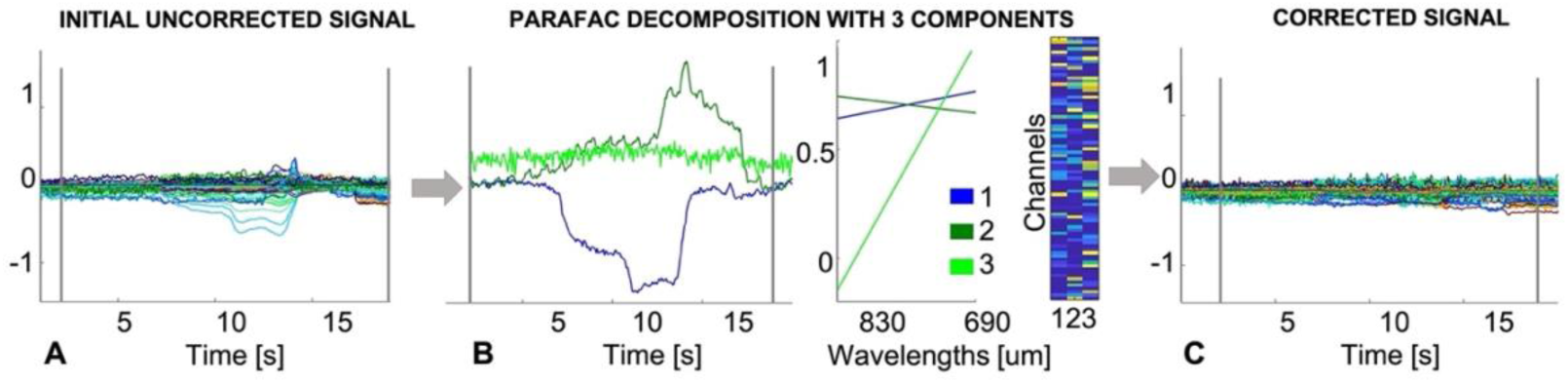
Example of target 3D PARAFAC decompositions to correct motion artifacts in a task-based fNIRS data set. Panel A Shows the initial uncorrected data segment. In panel B the temporal, spatial (channel) and wavelength (λ1|2 = 690|830 nm) signatures of the components identified with PARAFAC decomposition are presented, respectively. Panel C shows the corrected signal, illustrating the efficacy of movement artifact correction with PARAFAC after subtraction of two components (1 & 2). Y-axis is presented in arbitrary units.

Automatic and manual artifact detection agreed on a majority of the signal with an average concordance of 92.8%, so only minor manual adjustments had been applied. A total of 585 different artifactual events, on average 66 events per subject with a mean, minimal and maximal duration of 5, 0.7 and 21.2 s, respectively, were considered for correction. Comparison of the SNR included measures of every channel of both wavelengths. Repeated-measures ANOVA with Greenhouse-Geisser correction for the SNR of all conditions (Uncorrected, PARAFAC, tPCA and ICA) revealed a significant main effect, *F* (2.3, 44966.3) = 7550.7, *p* < 0.001, *ɲ* = 0.28. The uncorrected signal had the largest ratio of noise (−0.69), followed by the signal after ICA correction (−0.50), tPCA correction (−0.33), and PARAFAC correction (−0.31) (see Table 1). Pairwise contrasts ^52^ revealed that PARAFAC correction resulted in a significantly higher SNR compared to the uncorrected signal (*ɲ* = 0.45), and the signal after correction with ICA (*ɲ* = 0.13), both having a large effect size. Comparison of PARAFAC and tPCA, though statistically significant, revealed only a small effect size (*ɲ* = 0.002). Repeated-measures ANOVA with Greenhouse-Geisser correction for the Rλ revealed a significant difference between the five conditions: *F* (3.2, 30999.0) = 2686.46, *p* < 0.01, *ɲ* = 0.22. Mean Rλ coefficients showed the highest association between wavelengths for the uncorrected signal (0.74), followed by the signal after ICA correction (0.67), tPCA correction (0.53), PARAFAC correction (0.51) and the artifact-free reference signal (0.47) (see Table 1). Contrasts of Rλ coefficients between PARAFAC and ICA (*ɲ* = 0.14, large effect) as well as the uncorrected signal (*ɲ* = 0.35, large effect) suggested that wavelengths were greatly less correlated after PARAFAC correction. Correction with PARAFAC resulted in slightly higher Rλ coefficients compared to segments without artifacts (artifact-free reference, *ɲ* = 0.01, small effect), and slightly lower Rλ coefficients compared to correction with tPCA (*ɲ* = 0.01, small effect).

**Table 1.** Mean quality metrics for the evaluation of artifact correction in the task-based data set

| [Mean ± SD] | Artifact-free reference | PARAFAC | tPCA | ICA | Uncorrected artifact reference |
| --- | --- | --- | --- | --- | --- |
| Signal-to-noise ratio (dB) | N/A | $-0.31 \pm 0.58$ | $-0.33 \pm 0.57$ | $-0.50 \pm 0.64$ | $-0.69 \pm 0.65$ |
| Pearson's correlation between wavelengths ($R\lambda$) | $0.47 \pm 0.33$ | $0.51 \pm 0.38$ | $0.53 \pm 0.38$ | $0.67 \pm 0.38$ | $0.74 \pm 0.29$ |

Note. Results are displayed for the three decomposition techniques, parallel factor analysis (PARAFAC), target principal component analysis (tPCA) and independent component analysis (ICA), as well as for two reference signals, an artifact-free segment retrieved from the baseline before task onset, and the uncorrected signal consisting of all artifact segments before correction was applied.

### 3.2 Correction of simulated motion artifacts

As a second step, we evaluated the performance of PARAFAC to correct simulated artifacts with varying parameters, see Fig. 3 for the composition of the different scenarios. Three similarity indices (PRD, RMSE, R) for the agreement of the initial (NIRS_ini_) and the corrected signal (NIRS_PARAFAC/tPCA_), two quality measures (SNR, Rλ) and the reconstruction of the synthesized HRF were used to quantitatively compare performance of PARAFAC to tPCA. The sample for each index (PRD, RMSE, R, SNR) consisted of 164 measures, corresponding to channels of both wavelengths. Since Rλ is a correlation coefficient between both wavelengths, the sample consisted of 82 values. For each simulation condition (1a-e, 2a-f, 3a-d), indices were computed for three correction conditions (=within-subject factor), i.e. the signal corrected with PARAFAC or tPCA and the uncorrected signal.

Repeated measures ANOVA with Greenhouse-Geisser correction revealed statistically significant interactions for the type of correction and 1) the amplitude size (1a-e), PRD: *F* (8, 1630) = 67.14, *p* < 0.001, *ɲ*^*2*^_*partial*_ =0.248; RMSE: *F* (8, 1630) = 132.35, *p* < 0.001, *ɲ*^*2*^_*partial*_ =0.394; R: *F* (8, 1630) = 94.01, *p* < 0.001, *ɲ*^*2*^_*partial*_ = 0.316; SNR: *F* (8, 1630) = 214.90, *p* < 0.001, *ɲ*^*2*^_*partial*_ = 0.513; Rλ: *F* (8, 810) = 115.96, *p* < 0.001, *ɲ*^*2*^_*partial*_ = 0.534; 2) the onset delay of the second artifact (2a-f), PRD: *F* (10, 1956) = 4.98, *p* < 0.001, *ɲ*^*2*^_*partial*_ = 0.025; RMSE: *F* (10, 1956) = 8.93, *p* < 0.001, *ɲ*^*2*^_*partial*_ = 0.044; R: *F* (10, 1956) = 7.82, *p* < 0.001, *ɲ*^*2*^_*partial*_ = 0.038; SNR: *F* (10, 1956) = 7.18, *p* < 0.001, *ɲ*^*2*^_*partial*_ *=* 0.035; Rλ: *F* (10, 972) = 2.36, *p* = 0.01, *ɲ*^*2*^_*partial*_ = 0.024; and 3) the onset of the artifact relative to the HRF_sim_ (3a-d), PRD: *F* (6, 1304) = 26.67, *p* < 0.001, *ɲ*^*2*^_*partial*_ = 0.109; RMSE: *F* (6, 1304) = 52.74, *p* < 0.001, *ɲ*^*2*^_*partial*_ = 0.195; R: *F* (6, 1304) = 63.23, *p* < 0.001, *ɲ*^*2*^_*partial*_ = 0.225; SNR: *F* (6, 1304) = 61.31, *p* < 0.001, *ɲ*^*2*^_*partial*_ = 0.220; Rλ: *F* (6, 648) = 22.06, *p* = 0.009, *ɲ*^*2*^_*partial*_ = 0.170. Results of post-hoc comparisons with Tukey correction are illustrated in Fig. 6. using the example of PRD. The results of the other indices are mostly in line with those. Detailed results for the RMSE, R, SNR and Rλ can be found in the Supplemental Material in Fig. S2,Fig. **S3**, Fig. **S4**, andFig. **S5**, respectively.

**Fig. 6.**
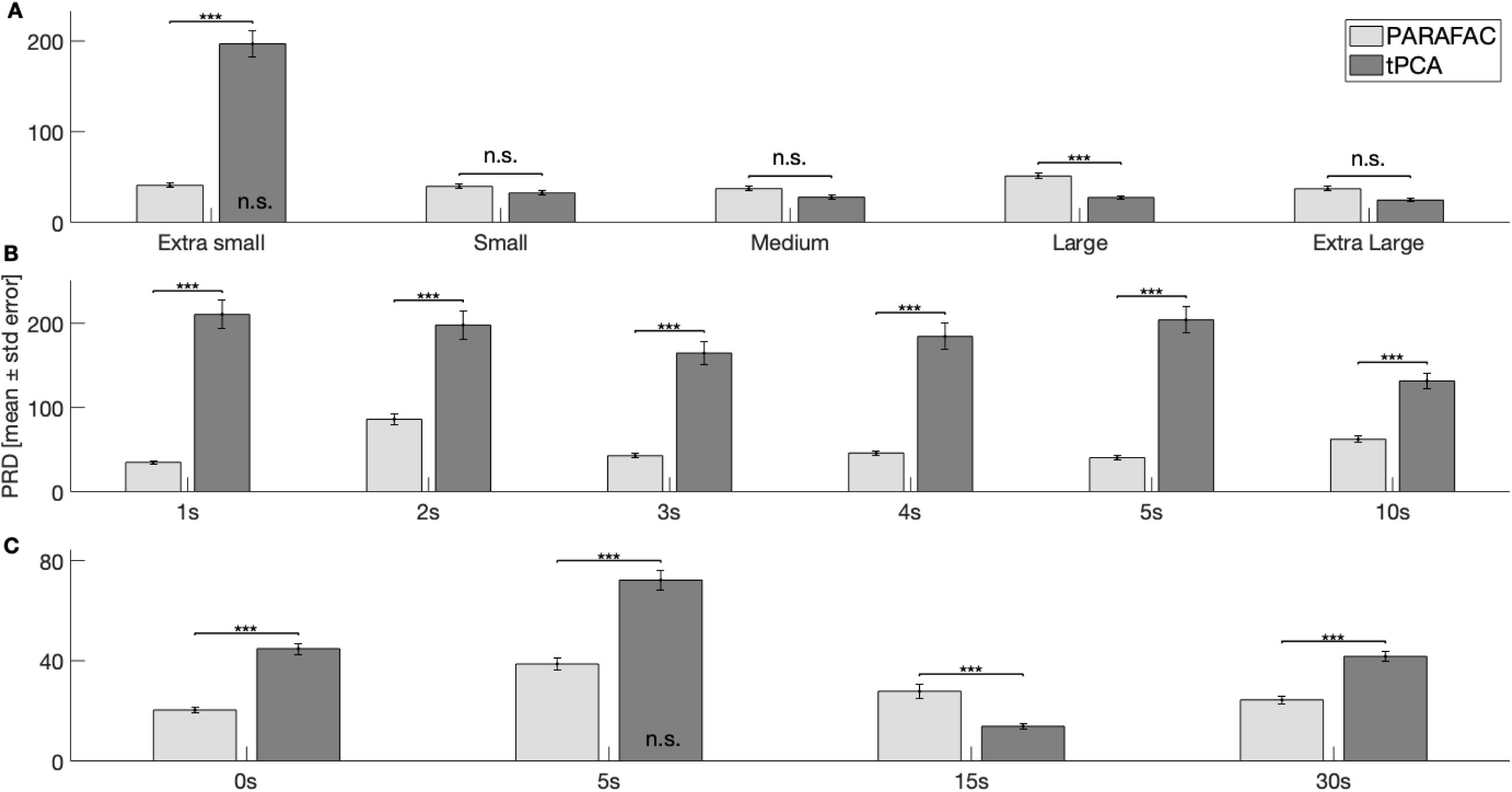
Evaluation of correction in simulated motion artifacts based on signal similarity. Simulations are identified on the x-axis. A: 1a-e) artifacts with different amplitude sizes; B: 2a-f): complex artifacts with two superimposed artifacts and an onset delay between the 1^st^ (A_1_) and 2^nd^ artifact (A_2_); and C: 3a-d) the onset of the artifact relative to the beginning of a simulated HRF. Performance of artifact correction is illustrated by the use of the similarity index percent root difference (PRD), where a lower value represents higher resemblance between the corrected and the initial clean fNIRS signal, hence a better correction of the artifact. Results are displayed separately for the correction with PARAFAC (light grey bars) and tPCA (grey bars). The PRD of the uncorrected signal is not displayed in this figure but differed significantly from both correction techniques in all conditions, except where specified (n.s.) otherwise inside the bar. Significance level are based on post-hoc tests with Tukey correction. *** p ≤ 0.001, n.s. p > 0.05. Uncorrected = NIRS_ini_ + artifact (A_1_) without correction, PARAFAC = NIRS_ini_ + artifact (A_1_/A_2_) after artifact correction with PARAFAC, tPCA = NIRS_ini_ + artifact (A_1_/A_2_) after artifact correction with tPCA.

Simulations 1a-e included artifacts with extra small, small, medium, large and extra-large amplitudes. Similarity indices (PRD, RMSE, R) revealed that both PARAFAC and tPCA resulted in a significant higher resemblance of the corrected signal and the initial signal compared to the uncorrected condition for all amplitude sizes except for the artifact with an extra small amplitude where tPCA did not lead to a significant improvement compared to the uncorrected signal. Similarly, both correction methods led to a significantly higher signal quality, i.e. lower SNR and Rλ, compared to the uncorrected signal with the same exception for tPCA in the extra small amplitude, where no difference was observed to the uncorrected signal. When artifacts had a small, medium or extra-large amplitude, correction methods obtained comparable results over all indices and did not statistically differ. The most consistent differences between correction methods can be observed for the artifact with extra small and large amplitudes. While similarity metrics and quality measures revealed that PARAFAC achieved significantly better results than tPCA for the correction of artifacts with an extra small amplitude, for artifacts with a large amplitude tPCA seemed to have a slight but statistically significant advantage as compared to PARAFAC.

Simulations 2a-f led to signals with different onset delays (1s, 2s, 3s, 4s, 5s, 10s) between two superimposed artifacts which allowed to create different complex artifacts. Over all conditions, notwithstanding their onset delay, similarity indices indicated a significantly higher overlap with the initial signal for the signal corrected with PARAFAC as compared to the one corrected with tPCA and the uncorrected signal. Similarly, quality metrics revealed a significantly higher signal quality after correction with PARAFAC compared to the correction with tPCA and the uncorrected signal. Nevertheless, tPCA also resulted in statistically significant better results as compared to the uncorrected signal in all conditions both with regard to the similarity metrics and the quality measures.

In simulations 3a-d the onset of an artifact varied relative to the beginning of a simulated HRF_sim_: +0s, +5s, +15s, +30s. Independent of the onset of the artifact, correction with PARAFAC resulted in a signal that showed a significantly higher overlap with the initial signal compared to the uncorrected condition. Similarly, quality measures showed a significant improvement of the signal’s quality after correction with PARAFAC compared to the uncorrected signal. tPCA achieved almost identical results as PARAFAC in comparison to the uncorrected signal, except when the artifact was placed shortly after the beginning of the HRF_sim_ (+5s) for which PRD and RMSE did not show a significant difference between the uncorrected signal and the one corrected with tPCA. Further, for artifacts at the very beginning (+0s), shortly after (+5s) or at the very end (+30s) of the HRF_sim_ (at the same time than the low-frequency increase or decrease that is intrinsic to the HRF_sim_), similarity indices revealed a significant better overlap of the corrected signal with the initial signal after correction with PARAFAC as opposed to tPCA. In contrast, tPCA compared to PARAFAC seemed to have a significant advantage for the correction of artifacts during the plateau of the HRF_sim_ (+15s), which might be explained by a higher orthogonality between the artifact and the signal at this onset. Quality measures however only partially support these findings, namely in favor of correction with tPCA for the artifact at the beginning (+0s) and in favor of PARAFAC for the artifact shortly after (+5s) the beginning of the HRF_sim_. When the artifact is at the very beginning of the HRF_sim_ (+0s), signal quality is however significantly higher after correction with tPCA as compared to PARAFAC. There is no statistically significant difference between signal quality for any of the correction methods when the artifact is placed at the very end of the HRF_sim_ (+30s). Further, results of the GLM revealed that in none of the conditions the simulated artifact led to an important reduction of the variance explained by the HRF_sim_ as compared to the initial signal (see Table 2). In line with the results of the similarity metrics, reconstruction of the HRF after correction with PARAFAC was qualitatively better as compared to tPCA when the artifact occurred at the beginning (+0s), shortly after (+5s) and at the end (+30s) of the HRF. In particular, for the artifact 5s after the onset of the HRF, reconstruction after correction with tPCA seemed disturbed. There was no important difference between correction methods for the reconstruction of the HRF when the artifact was during the plateau of the HRF (+15s).

**Table 2.**
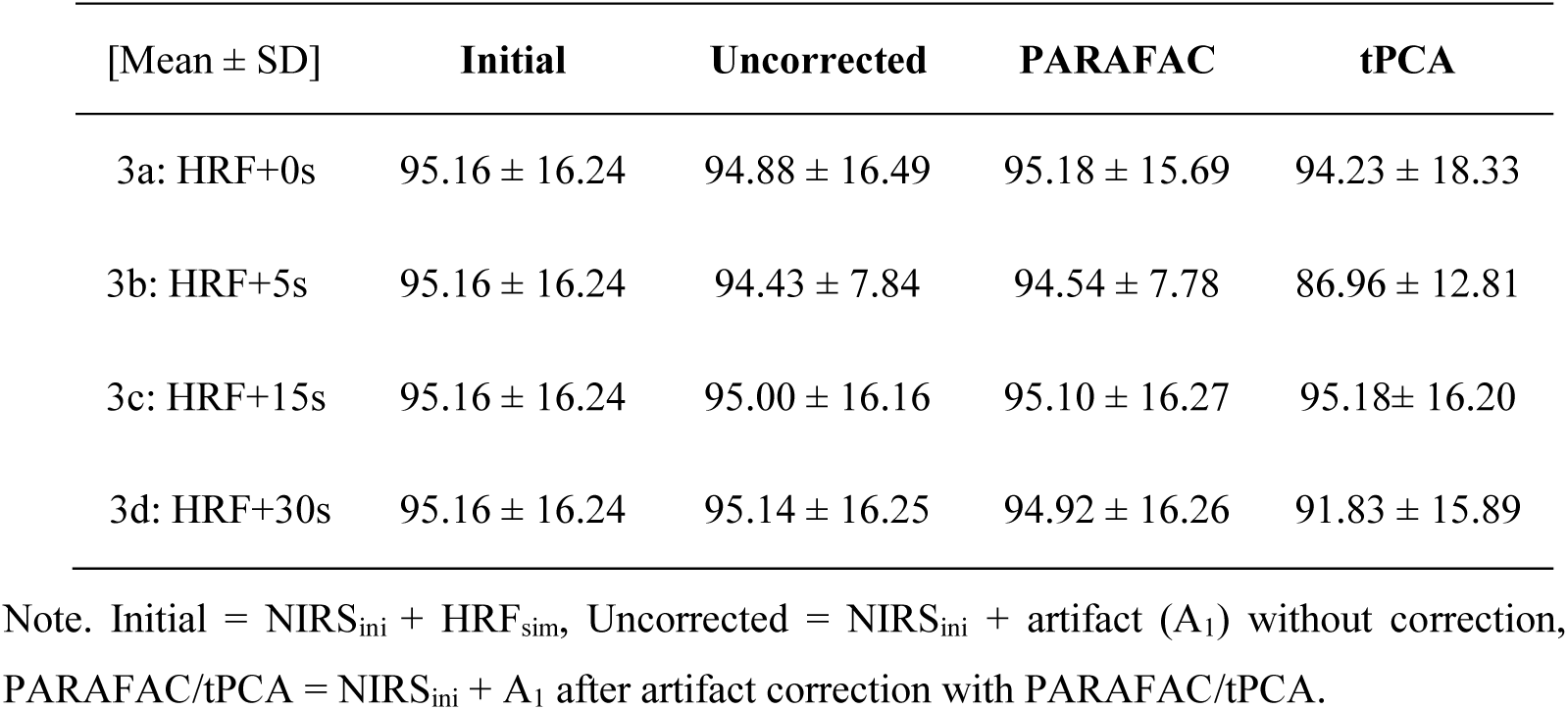
Percentage of explained variance by the HRF_sim_ based on the R^2^ of the general linear model

Orthogonality measure for the relation of the time courses of the raw signal and the artifact signal (Raw x A_1_: *r*_*a*_) were calculated for all simulations, orthogonality for the time courses of the raw signal with the first artifact and the second artifact (RawA_1_ x A_2_: *r*_*b*_) was only computed for simulations of complex artifacts (2a-f). A value of 1, represented maximum orthogonality between both signals meaning they are in a 90 degrees angle to each other. Any value smaller than 1 or even 0 indicated non-orthogonality meaning that they are in any other angle than 90 degrees, maximally 0 or 180 degrees to each other. Results of *r*_*a*_ revealed consistently high orthogonality for all simulations with varying amplitudes (Min = Max = 0.97) and with complex artifacts (Min = 0.97, Max = 0.98). Orthogonality for the simulations with the HRF revealed more variations (Min = 0.88, Max = 0.97) where the artifact 30s after the HRF_sim_ created the least orthogonal condition (Median ± standard error: 0.88 ± 0.03), followed by the simulation with the artifact at 0s (Median ± standard error: 0.93 ± 0.05), 5s (Median ± standard error: 0.96 ± 0.01) and 15s (Median ± standard error: 0.97 ± 0.01) of the HRF_sim_. Results of the second orthogonality measure *r*_*b*_ indicated that the onset delay of 2s led to the least orthogonal condition (Median ± standard error: 0.75 ± 0.07), followed by the delay of 3s (Median ± standard error: 0.93 ± 0.02) and 5s (Median ± standard error: 0.98 ± 0.02). When the second artifact had a delay of 1s, 4s or 10s, there was equally high orthogonality for all three conditions (Median ± standard error: 0.99 ± 0.02).

## 4 Discussion

Promising results have been reported by using parallel factor analysis (PARAFAC) for multidimensional (n ≥ 3) data analysis in EEG data ^13,28,42^. Since the fNIRS signal has inherently three dimensions, time × space × wavelength, we aimed to extend the application of PARAFAC to fNIRS and validate its use for artifact correction. First, we explored the usefulness of PARAFAC to correct movement artifacts in a data set of task-related fNIRS signals acquired during an expressive language paradigm. Performance of artifact correction was evaluated by the use of two signal quality metrics, (1) the SNR considering the signal’s variance during the simulated intervals after correction; and (2) the temporal similarity between both wavelengths during the simulated intervals after correction (Pearson’s correlation Rλ). Quality measures after correction with PARAFAC were compared to those obtained after the use of two commonly used decomposition methods in fNIRS (i.e. tPCA and ICA). Secondly, several scenarios with simulated artifacts in a clean resting-state signal were computed to assess the performance of artifact correction with PARAFAC and its homologue 2D target decomposition technique (tPCA) in a controlled setting. Simulated artifacts had five amplitude sizes, 6 levels of temporal overlap with a second artifact to create complex artifactual events and were added at four different time points of a simulated HRF. We compared the performance of both correction methods using (1) similarity indices describing the degree of correspondence between the corrected signal after removal of simulated artifacts and the artifact-free signal before simulation (RMSE, PRD, and R); and (2) the two quality measures, SNR and Rλ, already used for the task-related signal.

With regard to motion artifact correction in the task-based data set, the signal after artifact correction with PARAFAC had the smallest signal-to-noise ratio among the three methods (PARAFAC, tPCA, ICA), suggesting that more of the variance related to the artifact had been removed and a better signal quality was achieved. Similarly, correction with PARAFAC led to the lowest correlation between HbO and HbR indices, as well as the correlation index that most resembled the non-artifactual resting-state baseline reference. Even though the resting-state signal has been reported to show slightly different hemodynamic changes as compared to a stimuli-induced cerebral activity ^17,53^, we consider it a suitable reference for a time interval without artifacts, because it was not affected by articulation. In line with previous findings, our analyses also revealed a robust advantage in applying target corrections (tPCA and PARAFAC) instead of whole-block (ICA) correction, resulting in a better signal quality ^14,18^. ICA was indeed applied to the entire signal as reported in the literature ^17^, while tPCA and PARAFAC were used in a target manner, i.e. only decomposing the signal during specific intervals where artifacts had been detected. As proposed by Yücel and colleagues ^3^, target artifact correction prevents large changes in the overall composition of the signal, and exclusively corrects the noisy time interval of the artifact. It allows the decomposition to clearly sort a component representing the artifact signature, without considering the characteristics of these channels during the intervals without artifacts.

Target decomposition is thus beneficial for a precise identification of the artifact’s signatures and PARAFAC for artifact correction in fNIRS should also be applied in a target manner.

Results of the conducted simulations with controlled parameters suggest that artifact correction with PARAFAC led to a signal that corresponded more closely to the initial signal, as compared to the uncorrected signal. Similarly, PARAFAC yielded a significant better signal quality as compared to the uncorrected signal. This result is a preliminary validation of the use of PARAFAC for artifact correction in fNIRS signals and similarity measures show that the signal comes close to the original signal, thus it does not remove large parts of the physiological or relevant activity. While correction with tPCA in most cases also led to better results compared to the uncorrected signal, it has to be mentioned that its performance was less consistent over conditions. Further, in almost all scenarios correction with PARAFAC compared to tPCA led to either better or comparable results. Especially, when the artifact was simulated along with a simulated HRF, PARAFAC showed superior and more robust results compared to tPCA. According to the applied orthogonality measure, those conditions where tPCA showed a poor performance were exactly those where the artifact and the simulated HRF were less orthogonal. This is in line with the assumption that decomposition with PARAFAC is not affected by non-orthogonality. Only in two simulations, namely when the artifact had a large amplitude or was during the plateau of the simulated HRF (where the orthogonality between the HRF_sim_ and the artifact was the highest), correction with tPCA outperformed PARAFAC. Finally, results of the GLM indicate that artifact correction with PARAFAC did not affect the recovery of the simulated HRF, which suggests that it successfully targeted the artifact signature and did not induce changes of the signal that would alter interpretation of the underlying hemodynamic response. Even though this is an encouraging result, it has to be mentioned that similar results were achieved for the HRF_sim_ of the uncorrected signal, suggesting that the simulated artifact did not have a very strong impact on the signal and the interpretation of the HRF_sim_. This could be different for more complex or larger artifacts where correction may not sufficiently work. In most cases correction with tPCA allowed a similarly good recovery of the simulated HRF, but was strongly affected when an artifact was placed shortly after the beginning of the HRF_sim_, i.e. when the assumption of orthogonal artifact is not perfectly met. This is in line with the findings for the similarity and quality metrics for this scenario.

Regarding the comparison of both targeted decomposition techniques, the small effect size of the difference between tPCA and PARAFAC in the task-based data set suggests that both target correction techniques led to a similar reduction of the variance induced by motion artifacts and to an equal reduction of correlation between both wavelengths. Even though correction of simulated artifacts outlined certain advantages of PARAFAC compared to tPCA in some conditions, there was no consistent difference between both methods when applied to large data sets of real movement artifacts in a verbal fluency task. Since it was previously emphasized that performing accurate simulations of real motion artifacts is challenging, validation of an artifact correction technique in real data is a crucial step ^7^. We can conclude that its performance both in real and simulated data was generally equally good as tPCA and slightly better under certain conditions, suggesting the validation of PARAFAC as a new tool for artifact correction in fNIRS data.

What is more, PARAFAC’s core strength compared to tPCA is mainly related to its conceptualization. The use of a decomposition approach considering the multidimensional structure of the fNIRS signal where a unique decomposition is achieved with few constraints, i.e. without imposing orthogonality nor independence, is appropriate and seems advantageous compared to tPCA as well as ICA ^42,54^. Even though the results of the simulations provide some support that artifact correction with PARAFAC was not affected by non-orthogonality, orthogonality did not sufficiently vary among simulations and mostly reached a high orthogonality as we used resting state as background data. Nevertheless, given that in real data, the ground truth about the signal’s composition of noise is unknown and orthogonality cannot be verified, it is safe to say that PARAFAC represents a robust approach for artifact correction in fNIRS. Two-dimensional decomposition, such as ICA and tPCA, analyzes both wavelengths as independent measures of the fNIRS signal, even though they are in fact highly related, since they are acquired at the same location ^4^. Since the estimation of the hemodynamic signal is based on the signal of both wavelengths ^47^, reliable conclusions require the signal to be clean in both of them. It is also worth to notice that for artifact correction, we have followed in all our simulations the assumption that both wavelengths are affected by artifacts equally across channels (just with a different general scale). This is in fact the worst case for PARAFAC, as it will give better results if there are larger differences in all dimensions of the data. Therefore, we could expect an improved performance when artifacts have even small differences between the spatial distributions for the two wavelengths.

Decomposition with PARAFAC offers a rather easy way to make a selection of relevant components that differentiate between signatures related to artifacts or other relevant characteristics of the signal. Even if fNIRS is considered to be less sensitive to movements -as compared to other neuroimaging techniques such as fMRI and signal quality should be controlled during data acquisition as much as possible, appropriate tools for artifact correction are an essential part of preprocessing in fNIRS, particularly for data acquired in populations where cooperation is limited, such as children and clinical populations ^7,55,56^. PARAFAC thus appears to be a suitable and robust alternative to the currently used approaches and will be a valuable add-on for fNIRS studies and clinical examinations by minimizing the amount of non-usable data and improving data quality. That being said, the researcher always has to weigh the means of applying artifact correction and should not do so without visual inspection of the decomposition signature. It is the overall quality of a data set, i.e. the amount of clean signal, and the duration, extent and moment of the artifact that ought to influence the researcher’s choice to reject data sets, to apply artifact correction and to proceed with analysis.

### 4.1 Usefulness and limitations of PARAFAC

PARAFAC is a data-driven approach based on the linear relations of the three dimensions of the fNIRS signal ^23,26,28^. PARAFAC has the advantage of allowing for the three-way arrays of data to be uniquely decomposed into a sum of components, each of which is a trilinear combination of factors or signatures. The only statistical requirements of PARAFAC is that of a moderate linear independence across components, i.e. their time course, topography and wavelength characteristics. This is a less stringent requirement than previous models that underlie space/time decompositions (PCA or ICA). Each component provides characteristics of a particular pattern identified within the mixed measured fNIRS signal. When there are empirical or theoretical reasons to expect more than one relevant component (e.g. resting state functional connectivity analysis, epileptic activity, physiological aspects, etc.), PARAFAC would also allow disentangling several components. Moreover, when there are reasons to include other constraints such as orthogonality, non-negativity, smoothness and sparseness of the signatures, it has been shown that PARAFAC can also include them in the decomposition procedure ^23,57,58^.

Different from ICA, PARAFAC components can be ordered according to their importance in explaining the data’s variance ^59^. This is why many studies using ICA apply data reduction or clustering techniques such as PCA prior to the execution of ICA ^17,60^. Moreover, when using PARAFAC, the description of the data is based on more dimensions, and with less theoretical constraints (orthogonality, independence) which lead to the identification of fewer relevant components ^13,28,61,62^. Compared to ICA, the selection of the relevant components is thus simplified. Importantly, when using PARAFAC, it is not recommended to ask for more than five components, as this increases the risk of overfitting the data with the decomposition model. Usually, three to five components sufficiently describe the signal’s relevant characteristics ^29^. Similar to PCA, a typical choice made when using PARAFAC analysis, is that of ordering the extracted components according to their contribution to explaining the variance of the data. In this sense, it can be expected that the first PARAFAC component will always represent the highest activity, which, in the case of target artifact correction, will correspond to the artifact activity. This makes PARAFAC also a promising choice to explore the development of simple automatic methodologies for the detection and correction of such artifacts and future studies should be devoted to this aim.

Some challenges of PARAFAC have already been discussed in previous studies ^13,28^. One limitation is related to PARAFAC’s assumption of linear relations of the temporal characteristics between different channels (which also applies to (t)PCA and ICA). Despite being one of the most common and simple models, imposing linearity may not entirely reflect complex cerebral processes ^63^. Some attempts have been made to introduce approaches that tolerate non-linear relations ^64,65^. However, to date, linear models remain the most popular and adequate approach for analysis of macroscopic data in neuroscience. Thus, it represents a general limit of the domain, rather than one specific to PARAFAC. Another potential limitation of our implementation of PARAFAC might arise from the specific use of the simple alternating least-squares algorithm to perform the decomposition. Other methods have been proposed for estimating PARAFAC models, but some studies have shown that, given the uniqueness of the solution, they do not usually outperform the simple least-squares technique ^66^.

The need for careful preprocessing of data before characterizing brain activity constitutes another limitation. When PARAFAC was introduced for the analysis of EEG data, the authors emphasized the importance of searching for constant factors, outliers, and degeneracy ^13,27^. Detailed information on how to deal with these aspects can be found in previous literature ^13^. In the current study, the implementation of PARAFAC into the LIONirs toolbox ^35^, gave us a certain flexibility regarding the channels and time segments to be included. Thus, we were able to exclude deviant channels prior to use PARAFAC.

Finally, among the applications of PARAFAC not yet tested in fNIRS, is its use to identify patterns of brain activity in the hemodynamic response as it has been applied in EEG data analysis ^13,29,42^. Compared to other techniques such as GLM or global averaging who treat each channel independently and provide a narrower spatial representation of the dominant activation, PARAFAC analysis could reveal a wider distributed activation pattern. This would strongly correspond to the current understanding of cerebral processing, whereby mostly large-scale networks, and not isolated regions, are considered to be involved in various cognitive processes ^55,67,68^. PARAFAC could thus appear to be suitable for reflecting cerebral processes occurring in distributed networks, rather than for the identification of a specific core region. This is obviously given a sufficient fNIRS covering. Moreover, the topographic signature would correspond to a whole time course of the extracted activations and can easily be subjected to diffusion optical tomography in order to locate the HbO and HbR concentration changes in the brain cortex ^69^. Another potential application is using PARAFAC as a screening tool. For instance, Miwakeichi and colleagues ^13^ applied PARAFAC to an EEG dataset in order to extract one component related to ocular movement artifacts, and subsequently used the PARAFAC analysis fixing spatial and spectral signatures of that component to screen a second dataset in order to identify and correct similar artifacts. This application can also support the development of a detection method as has already be done in other fields such as detection of epileptic seizures ^29,70^ and for brain-computer interfacing ^71–74^. Although this can be useful for correcting artifacts that have consistent topographical and wavelength profiles, movement artifacts were mostly related to articulation in our study, often showing quite different signatures. A different paradigm may enable the testing of this application of PARAFAC in fNIRS.

Our findings also encourage the general application of PARAFAC for multidimensional decomposition in neuroimaging, where data can often be described in more than two dimensions. Beyond the obvious dimensions of a technique such as that of time × space × wavelength (fNIRS) or frequency (EEG), characteristics of the task paradigm or group variables could also be considered in the decomposition model ^42^. For instance, if a “group” dimension was to be included, groups could be compared regarding their relative weight over the main cerebral activation component that would have been outlined from the PARAFAC model.

### 4.2 Limits of the current study

The use of a language paradigm allowed for the induction of artifacts that were mainly related to articulation, which limits the generalization from our results. From video recordings, however, we were able to acknowledge that swallowing, jaw or tongue movements, eye blinks, and frowning also induced artifacts in our dataset. Even though the simulated artifacts were also based on one identified during the task-condition, parameters were modified and allowed to create noise with different characteristics. Further fNIRS studies using PARAFAC, and including different task paradigms, resting-state, and recording in naturalistic environments when participants are moving, could show its utility for a wider range of artifacts (e.g. baseline shifts or physiology).

The validation of PARAFAC for artifact correction was conducted by comparing results with two other decomposition techniques. This could be considered one limit of the present study, because we did not consider other correction approaches for comparison. The objective of this study was not to conduct exhaustive and systematic comparisons in order to identify the best method for artifact correction, but rather to introduce PARAFAC as an alternative and adequate method in fNIRS analysis. Future studies could therefore expand the validation of PARAFAC and extend comparisons of its performance to other tools that do not necessarily use decomposition, such as spline interpolation or wavelet filtering. With relation to that, even if there were to be a slight advantage of PARAFAC over tPCA in artifact correction results, the small difference might not have a concrete impact on the applicability of those two methods. Qualitative differences between tPCA and PARAFAC regarding the type of artifacts and corrected signal should be further assessed and described in subsequent studies to better understand how their efficacy may vary according to the artifact’s characteristics.

Last but not least, PARAFAC’s components related to artifacts were selected during a visual inspection based on their temporal overlap with the artifact, i.e. sudden change of amplitude, and those showing similar weights for both wavelengths. The smallest number of possible components that sufficiently corrected the artifact were selected. Even though our results show that this procedure led to satisfying correction, it is not a fully standardized procedure which may hamper an easier, automatic correction that would reduce the need of the researcher’s expertise in detecting and adequately applying correction. This issue could be addressed using the screening procedure mentioned in the previous section or using other templates for the artifacts of interest or by applying the usual automatic rules based on amplitude characteristics to the temporal signatures obtained by PARAFAC instead of the original mixed data. The goal of the current study was not to propose a method for artifact detection but to validate the relative efficacy of these methods in artifact correction. It would be interesting for future studies to address the development of a more automatic procedure based on PARAFAC analysis contributing to a faster and standardized processing pipeline.

## 5 Conclusions

PARAFAC has the advantage to simultaneously treat both wavelengths or HbO and HbR during fNIRS data analyses. Our findings from real task-related signals and controlled simulations validate previous results in EEG data and promote multidimensional decomposition with PARAFAC as a promising new tool for the correction of movement artifacts in fNIRS. Precisely, PARAFAC achieves comparable results as tPCA when the artifact’s signature is simple and clearly distinguishable from the signal. It outperforms tPCA mainly when artifacts have small amplitude, show a complex temporal signature or when they co-occur during an HRF. These results can partially be attributed to low orthogonality between signal and noise. The advantages of PARAFAC and tPCA, as compared to ICA, are consistent and seem to be due to the target application. Further, PARAFAC has a strong advantage compared to both 2D decomposition techniques because it offers a unique decomposition without orthogonality or independence constraints, hence represents a robust decomposition technique. The validation of its use paves the way for future use in fNIRS research to extract relevant signatures represented in the fNIRS signal.

## 6 Supplemental Material

**Fig. S1.**
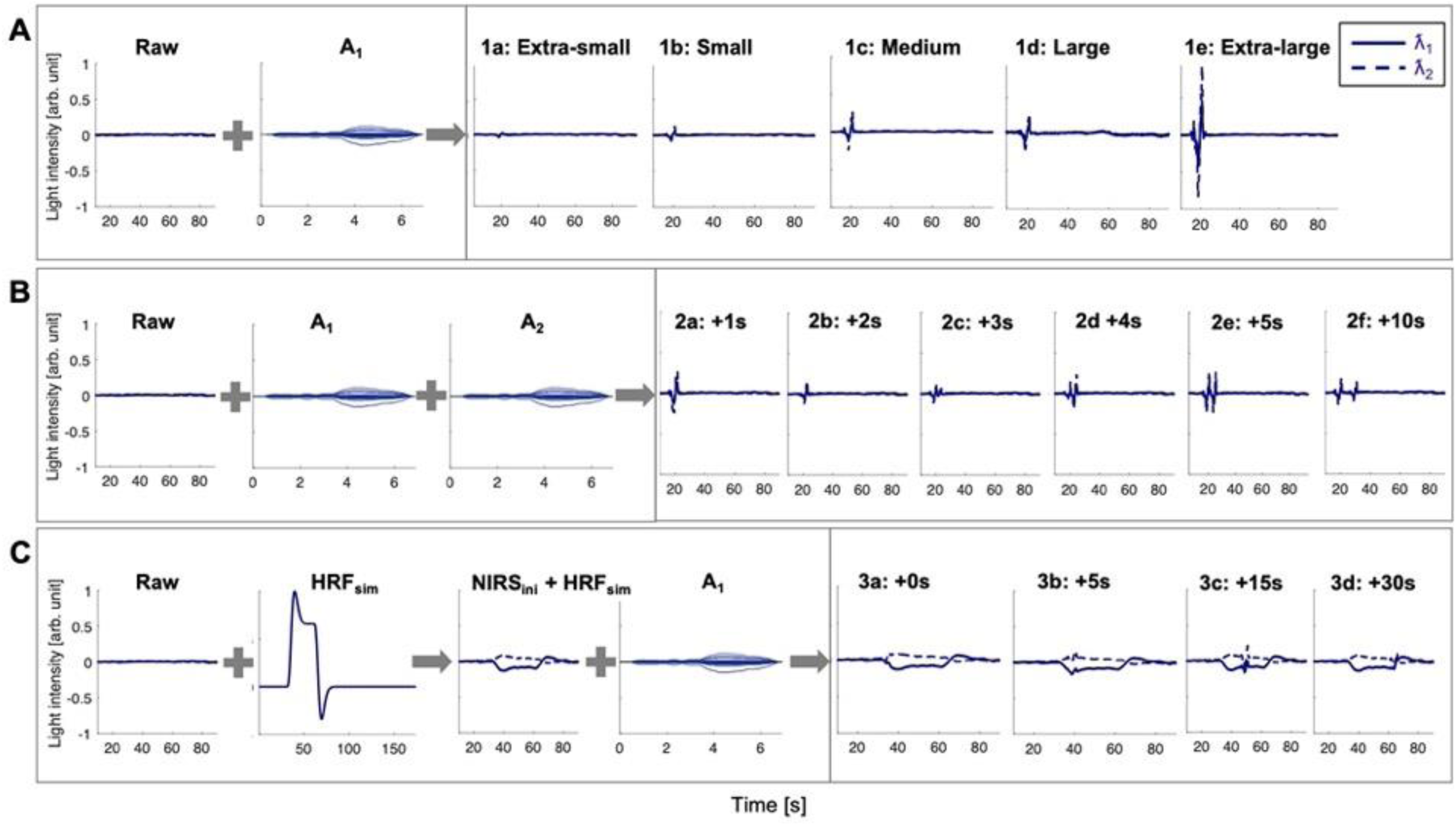
Parameters of all simulations. A real motion artifact (A_1/2_) and a synthesized HRF (HRF_sim_) were added to a normalized resting-state fNIRS signal (Raw). A shows simulations 1a-e with an artifact of varying amplitude sizes, B simulations 2a-f with a complex artifact where the onset of the second artifact (A_2_) was varied relative to the onset of the first artifact (A_1_), and C simulations 3a-d with varying onset of the artifact relative to the beginning of the HRF_sim_. λ1|2 = 690|830 nm.

**Fig. S2.**
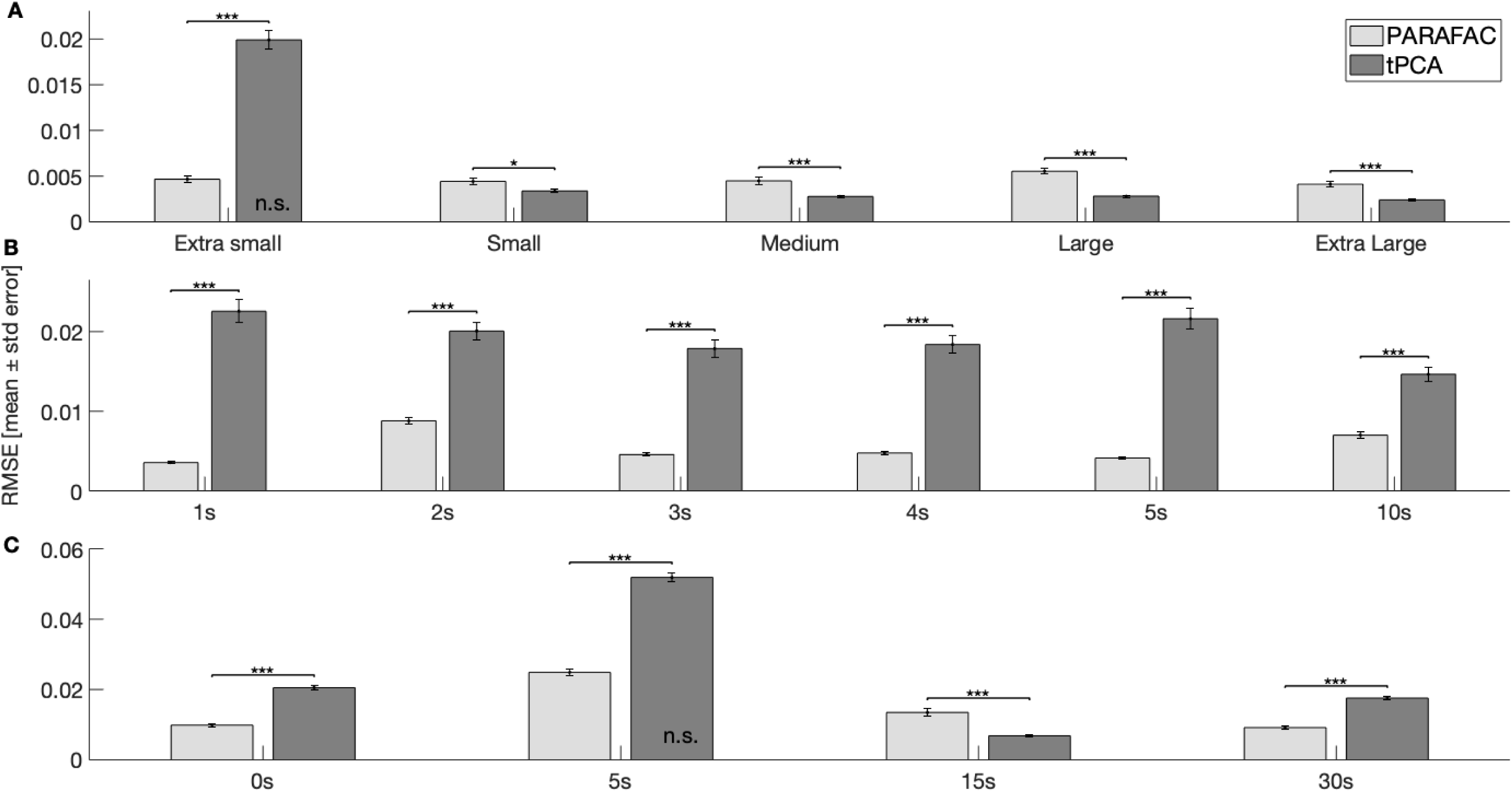
Evaluation of correction in simulated motion artifacts based on signal similarity by the use of the root mean square error (RMSE). Simulations are identified on the x-axis. A: 1a-e) artifacts with different amplitude sizes; B: 2a-f): complex artifacts with two superimposed artifacts and an onset delay between the 1^st^ (A_1_) and 2^nd^ artifact (A_2_); and C: 3a-d) the onset of the artifact relative to the beginning of a simulated HRF. A lower RMSE represents higher resemblance between the corrected and the initial clean fNIRS signal, hence a better correction of the artifact. Results are displayed separately for the correction with PARAFAC (light grey bars) and tPCA (grey bars). The RMSE of the uncorrected signal is not displayed in this figure but differed significantly from both correction techniques in all conditions, except where specified otherwise inside the bar. Significance level are based on post-hoc tests with Tukey correction. * p ≤ 0.05, *** p ≤ 0.001, n.s. p > 0.05. Uncorrected = NIRS_ini_ + artifact (A_1_) without correction, PARAFAC = NIRS_ini_ + artifact (A_1_/A_2_) after artifact correction with PARAFAC, tPCA = NIRS_ini_ + artifact (A_1_/A_2_) after artifact correction with tPCA.

**Fig. S3.**
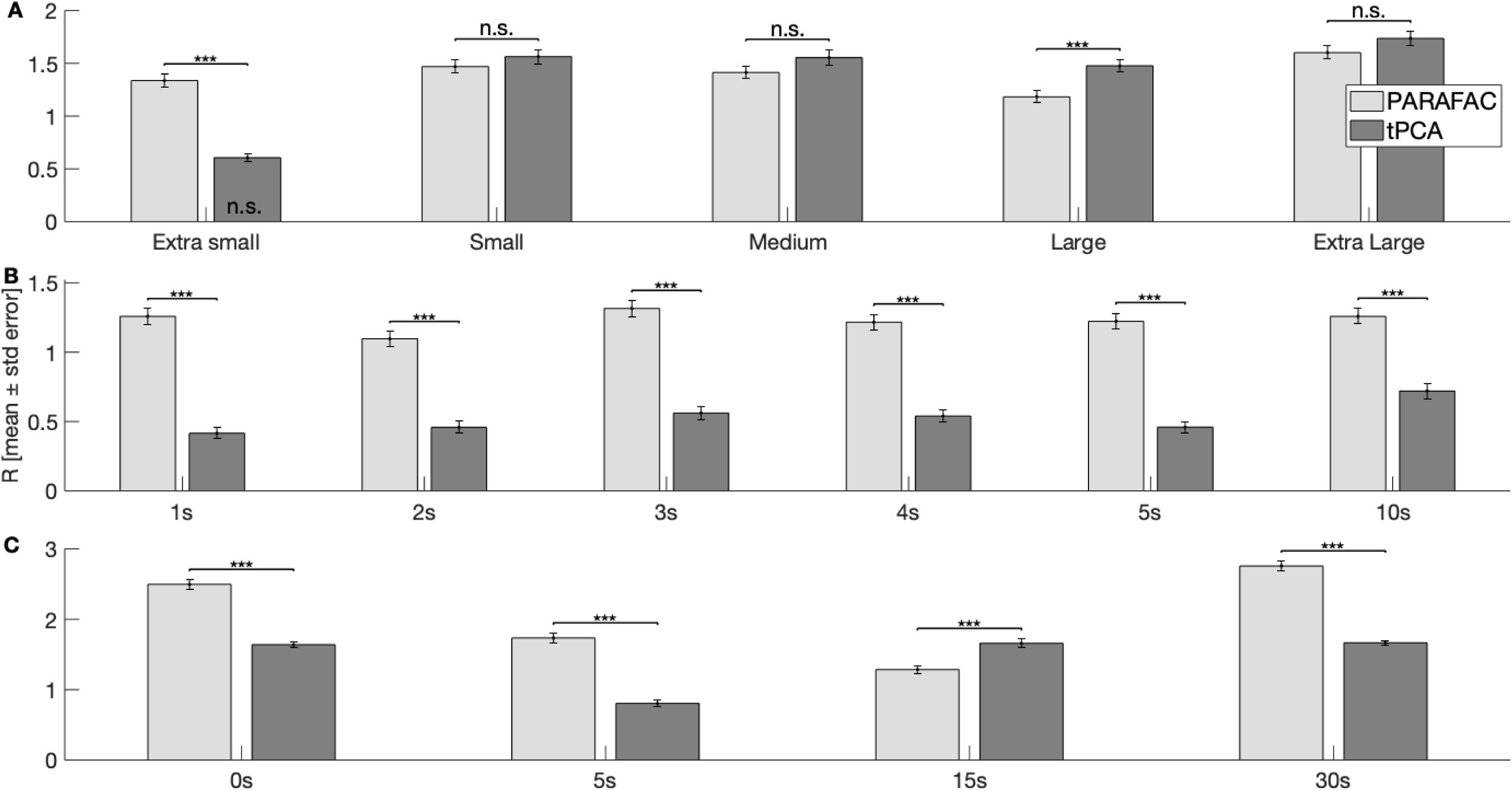
Evaluation of correction in simulated motion artifacts based on signal similarity by the use of the Pearson product-moment correlation coefficient (R). Simulations are identified on the x-axis. A: 1a-e) artifacts with different amplitude sizes; B: 2a-f): complex artifacts with two superimposed artifacts and an onset delay between the 1^st^ (A_1_) and 2^nd^ artifact (A_2_); and C: 3a-d) the onset of the artifact relative to the beginning of a simulated HRF. A higher R represents higher resemblance between the corrected and the initial clean fNIRS signal, hence a better correction of the artifact. Results are displayed separately for the correction with PARAFAC (light grey bars) and tPCA (grey bars). The R of the uncorrected signal is not displayed in this figure but differed significantly from both correction techniques in all conditions, except where specified otherwise inside the bar. Significance level are based on post-hoc tests with Tukey correction. *** p ≤ 0.001, n.s. p > 0.05. Uncorrected = NIRS_ini_ + artifact (A_1_) without correction, PARAFAC = NIRS_ini_ + artifact (A_1_/A_2_) after artifact correction with PARAFAC, tPCA = NIRS_ini_ + artifact (A_1_/A_2_) after artifact correction with tPCA.

**Fig. S4.**
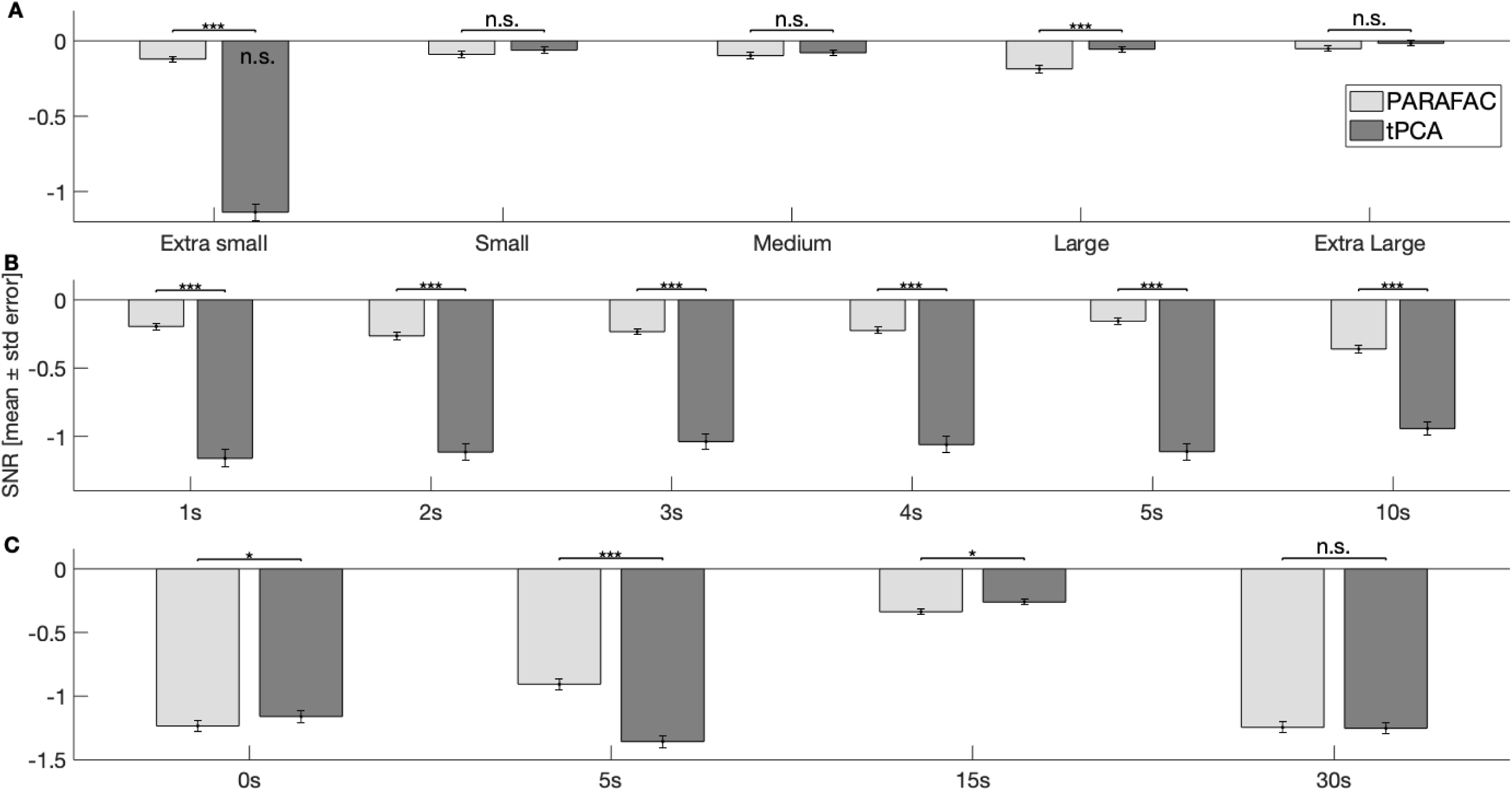
Evaluation of correction in simulated motion artifacts based on signal quality by the use of the signal-to-noise ratio (SNR). Simulations are identified on the x-axis. A: 1a-e) artifacts with different amplitude sizes; B: 2a-f): complex artifacts with two superimposed artifacts and an onset delay between the 1^st^ (A_1_) and 2^nd^ artifact (A_2_); and C: 3a-d) the onset of the artifact relative to the beginning of a simulated HRF. An SNR closer to 0 represents a more equal ratio between the signal’s variation during the artifact period and the clean signal. Results are displayed separately for the correction with PARAFAC (light grey bars) and tPCA (grey bars). The SNR of the uncorrected signal is not displayed in this figure but differed significantly from both correction techniques in all conditions, except where specified otherwise inside the bar. Significance level are based on post-hoc tests with Tukey correction. * p ≤ 0.05, *** p ≤ 0.001, n.s. p > 0.05. Uncorrected = NIRS_ini_ + artifact (A_1_) without correction, PARAFAC = NIRS_ini_ + artifact (A_1_/A_2_) after artifact correction with PARAFAC, tPCA = NIRS_ini_ + artifact (A_1_/A_2_) after artifact correction with tPCA.

**Fig. S5.**
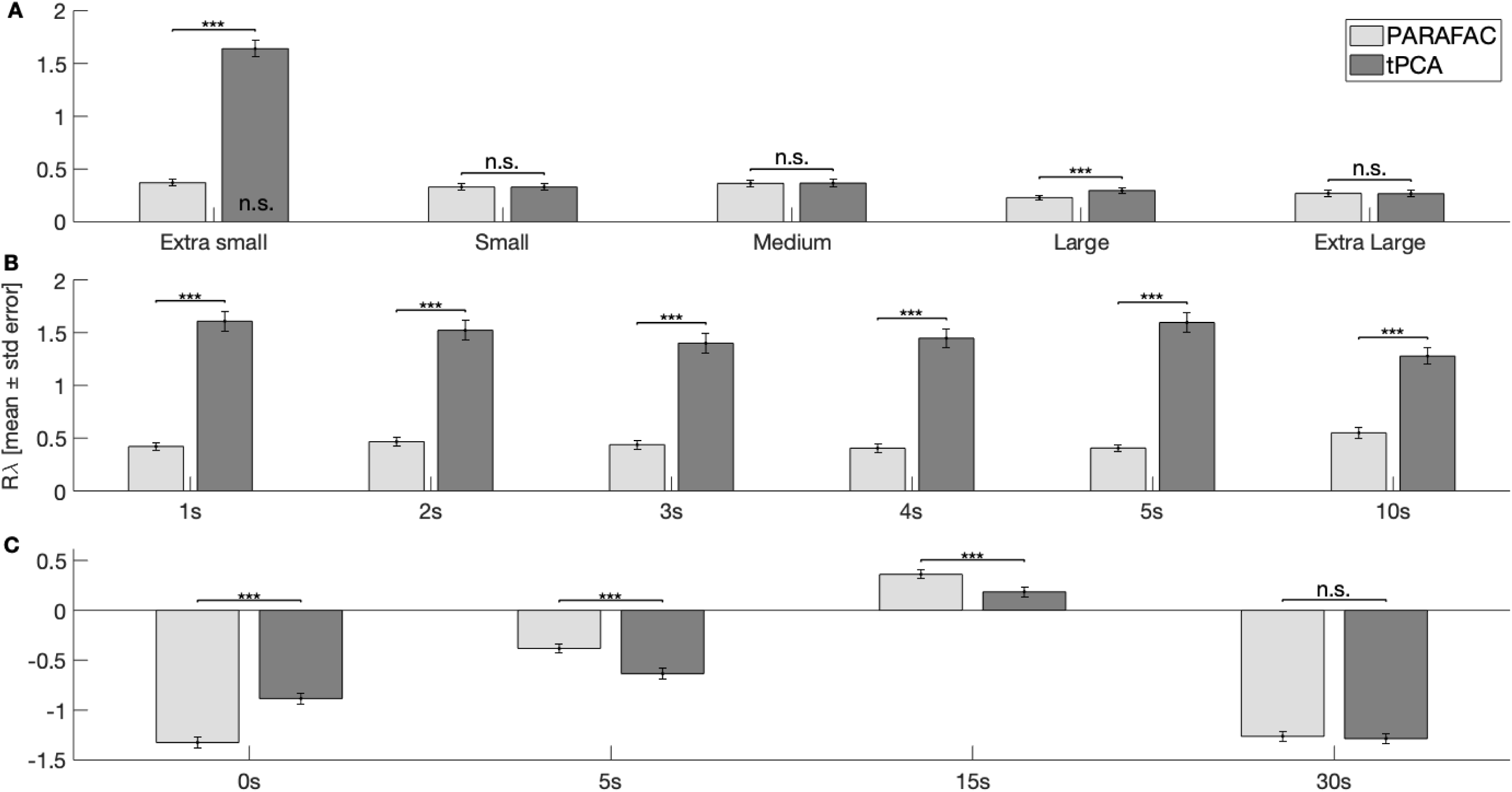
Evaluation of correction in simulated motion artifacts based on signal quality by the use of the Pearson’s correlation between wavelengths (Rλ). Simulations are identified on the x-axis. A: 1a-e) artifacts with different amplitude sizes; B: 2a-f): complex artifacts with two superimposed artifacts and an onset delay between the 1^st^ (A_1_) and 2^nd^ artifact (A_2_); and C: 3a-d) the onset of the artifact relative to the beginning of a simulated HRF. Rλ in a clean signal is usually lower than in artifacted signals. Results are displayed separately for the correction with PARAFAC (light grey bars) and tPCA (grey bars). The Rλ of the uncorrected signal is not displayed in this figure but differed significantly from both correction techniques in all conditions, except where specified otherwise inside the bar. Significance level are based on post-hoc tests with Tukey correction. *** p ≤ 0.001, n.s. p > 0.05. Uncorrected = NIRS_ini_ + artifact (A_1_) without correction, PARAFAC = NIRS_ini_ + artifact (A_1_/A_2_) after artifact correction with PARAFAC, tPCA = NIRS_ini_ + artifact (A_1_/A_2_) after artifact correction with tPCA.

## 8 Acknowledgments/Funding sources

The authors warmly thank all participants of the study, as well as Catherine Bernard, Kathya Martel and Solène Fourdain for their support during data acquisition.

This work was funded by the National Science and Engineering Research Council of Canada (NSERC) (#2015-04199 and # CGSD3-518503-2018); the Canada Research Chairs (#950-232661); the Ministère des Affaires internationales et de la Francophonie du Québec; the Fonds de Recherche du Québec Santé (FRQS, #28811 and #35450); the Fonds de recherche du Québec – Nature et technologies (FRQNT, #255473); the SickKids Foundation (#NI16-058); the Quebec Bio-Imaging Network (QBIN, FRQS - Réseaux de recherche thématique, #35450); the Savoy Foundation, and Graduate and Postdoctoral Studies Faculty (ESP) of the Université de Montréal.

---

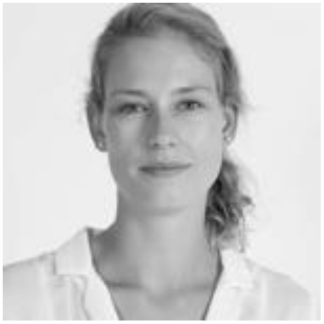

**Alejandra M. Hüsser** is a Ph.D. candidate in neuropsychology at the Université de Montréal. She has received a B.Sc. and M.Sc. degree in psychology with a specialization in cognitive sciences and neuropsychology from the University of Zurich in 2013 and 2015, respectively. Her research interests include childhood neurodevelopment, neurodevelopmental pathologies, epilepsy, cerebral language networks, language functions, neuroimaging and imaging analysis. In 2020, Alejandra was awarded a prestigious studentship by the Savoy Foundation.

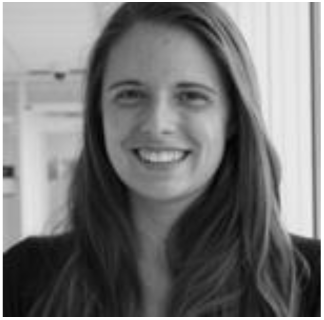

**Laura Caron-Desrochers** is a doctorate candidate in clinical neuropsychology at the Université de Montréal. She has received a B.Sc. degree in Psychology from the same university in 2015. Her research interests include the application of fNIRS to study early cerebral neurodevelopment and the influence of prenatal environment on the establishment of cerebral language networks.

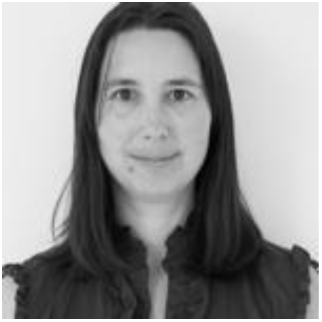

**Julie Tremblay** is a biomedical engineer in the LIONlab. She graduated with a B.Sc. degree in electrical engineering from the Université de Sherbrooke in 2006. Since, she specialized in data analysis, signal processing, and software development for neuroimaging data. She has participated in several projects using a wide range of imaging techniques such as NIRS, EEG, iEEG, MEG, DTI, and fMRI. Her research interests are in the integration of algorithms and understanding brain development.

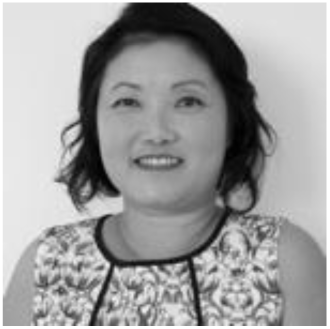

**Phetsamone Vannasing** joined CHU Sainte-Justine in 1993 as a research coordinator and electrophysiologist. Her task is to coordinate research protocols and teach students the technical aspects of data recording, acquisition, and analysis. She set up a state-of-the-art multimodal recording platform combining EEG, NIRS, eye-tracking, and electro-dermal systems. Her research interests include neurodevelopment in clinical and healthy populations. She received the Research Professionals Excellence awards of the Fonds de recherche du Québec-Santé and CHU Sainte-Justine.

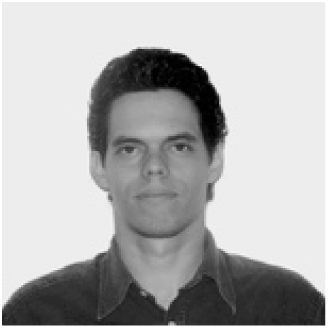

**Eduardo Martínez-Montes**, Ph.D., is a Senior Researcher at the Cuban Center for Neurosciences, and the director of the Human Brain Mapping Division. He has awarded with the Medal Carlos J. Finlay, the highest order given by the Cuban government for outstanding contributions to science. He has contributed to develop neuroscientific analysis, including multidimensional space-time-frequency analysis in EEG, multimodal EEG-fMRI analysis and estimation of electrophysiological brain sources. His current research focuses on the development of statistical models and the implementation of neuroinformatics tools for the analysis of brain activity.

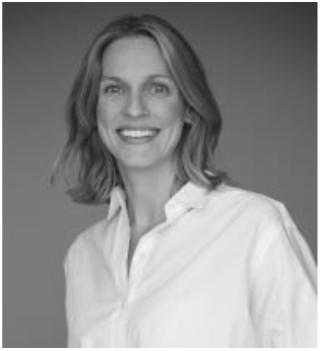

**Anne Gallagher**, Ph.D. and neuropsychologist, holds a Canada Research Chair in Child Neuropsychology and Brain Imaging. She is an Associate Professor at the Department of Psychology of the Université de Montréal and the Director of the LIONlab. She uses NIRS, EEG, and neuropsychological assessment to identify predictive brain markers of neurodevelopmental outcomes in children with various clinical conditions. She is a pioneer in the use of NIRS-EEG in clinical settings, notably developing presurgical evaluation techniques in patients with epilepsy.

## References

1. F. F. Jobsis, “Noninvasive, infrared monitoring of cerebral and myocardial oxygen sufficiency and circulatory parameters,” Science 198(4323), 1264–1267, American Association for the Advancement of Science (1977) [doi:10.1126/SCIENCE.929199].

2. S. Lloyd-Fox, A. Blasi, and C. E. Elwell, “Illuminating the developing brain: The past, present and future of functional near infrared spectroscopy,” Neuroscience and Behavioural Reviews 34, 269–284 (2010) [doi:10.1016/j.neubiorev.2009.07.008].

3. M. A. Yücel et al., “Targeted principle component analysis: A new motion artefact correction approach for near-infrared spectroscopy,” Journal of Innovative Optical Health Sciences 07(2), 1350066 (2014) [doi:10.1142/S1793545813500661].

4. X. Cui, S. Bray, and A. L. Reiss, “Functional near infrared spectroscopy (NIRS) signal improvement based on negative correlation between oxygenated and deoxygenated hemoglobin dynamics,” Neuroimage 49(4), 3039–3046 (2010) [doi:10.1016/j.neuroimage.2009.11.050].

5. R. J. Cooper et al., “A systematic comparison of motion artifact correction techniques for functional near-infrared spectroscopy,” Frontiers in Neuroscience 6(OCT), 1–10 (2012) [doi:10.3389/fnins.2012.00147].

6. S. Tak and J. C. Ye, “Statistical analysis of fNIRS data: A comprehensive review,” Neuroimage 85, 72–91 (2014) [doi:10.1016/j.neuroimage.2013.06.016].

7. S. Brigadoi et al., “Motion artifacts in functional near-infrared spectroscopy: A comparison of motion correction techniques applied to real cognitive data,” Neuroimage 85, 181–191 (2014) [doi:10.1016/j.neuroimage.2013.04.082].

8. H. Zhang et al., “Is resting-state functional connectivity revealed by functional near-infrared spectroscopy test-retest reliable?,” Journal of Biomedical Optics 16(6), 067008 (2011) [doi:10.1117/1.3591020].

9. G. James et al., “Classification,” in An Introduction to Statistical Learning, pp. 127–173, Springer, New York, NY (2013) [doi:10.1007/978-1-4614-7138-7_4].

10. X. Zhang et al., “Signal processing of functional NIRS data acquired during overt speaking,” Neurophotonics 4(04), 1 (2017) [doi:10.1117/1.NPh.4.4.041409].

11. K. Peng et al., “fNIRS-EEG study of focal interictal epileptiform discharges,” Epilepsy Research 108, 491–505 (2014) [doi:10.1016/j.eplepsyres.2013.12.011].

12. H. Becker et al., “Multi-way space-time-wave-vector analysis for EEG source separation,” Signal Processing 92, 1021–1031 (2011) [doi:10.1016/j.sigpro.2011.10.014].

13. F. Miwakeichi et al., “Decomposing EEG data into space–time–frequency components using Parallel Factor Analysis,” Neuroimage 22(3), 1035–1045 (2004) [doi:10.1016/j.neuroimage.2004.03.039].

14. H. F. Behrendt et al., “Motion correction for infant functional near-infrared spectroscopy with an application to live interaction data,” Neurophotonics 5(01), 1 (2018) [doi:10.1117/1.NPh.5.1.015004].

15. S. Jahani et al., “Motion artifact detection and correction in functional near-infrared spectroscopy: A new hybrid method based on spline interpolation method and Savitzky-Golay filtering,” Neurophotonics 5(1), 1, SPIE-Intl Soc Optical Eng (2018) [doi:10.1117/1.NPh.5.1.015003].

16. I. Tachtsidis and F. Scholkmann, “False positives and false negatives in functional near-infrared spectroscopy: issues, challenges, and the way forward,” Neurophotonics 3(3), 031405 (2016) [doi:10.1117/1.NPh.3.3.031405].

17. H. Zhang et al., “Functional connectivity as revealed by independent component analysis of resting-state fNIRS measurements,” Neuroimage 51, 1150–1161 (2010) [doi:10.1016/j.neuroimage.2010.02.080].

18. M. A. Kamran, M. M. N. Mannan, and M. Y. Jeong, “Cortical Signal Analysis and Advances in Functional Near-Infrared Spectroscopy Signal: A Review,” Frontiers in Human Neuroscience 10, 261 (2016) [doi:10.3389/fnhum.2016.00261].

19. A. Jung et al., “Fastgeo-a histogram based approach to linear geometric ICA,” in Proceedings of ICA Vol. 1, pp. 349–354 (2001).

20. P. Comon, “Independent component analysis, A new concept?,” Signal Processing 36(3), 287–314 (1994) [doi:10.1016/0165-1684(94)90029-9].

21. C. Jutten and J. Herault, “Blind separation of sources, part I: An adaptive algorithm based on neuromimetic architecture,” Signal Processing 24(1), 1–10 (1991) [doi:10.1016/0165-1684(91)90079-X].

22. R. A. Harshman, “Foundations of the PARAFAC procedure: Models and conditions for an ‘explanatory’ multimodal factor analysis,” in UCLA Working Papers in Phonetics 16, University Microfilms, Ann Arbor, Michigan (1970).

23. R. Bro, “Multi-way Analysis in the Food Industry Models, Algorithms, and Applications,” Royal Veterinary and Agricultural University (1998).

24. J. D. Carroll and J.-J. Chang, “Analysis of individual differences in multidimensional scaling via an n-way generalization of ‘Eckart-Young’ decomposition,” Psychometrika 35(3), 283–319 (1970) [doi:10.1007/BF02310791].

25. L. R. Tucker, “Some mathematical notes on three-mode factor analysis,” Psychometrika 31(3), 279–311 (1966).

26. J. Möcks, “Decomposing event-related potentials: A new topographic components model,” Biological Psychology 26(1–3), 199–215 (1988) [doi:10.1016/0301-0511(88)90020-8].

27. A. S. Field and D. Graupe, “Topographic component (Parallel Factor) analysis of multichannel evoked potentials: Practical issues in trilinear spatiotemporal decomposition,” Brain Topography 3(4), 407–423 (1991) [doi:10.1007/BF01129000].

28. E. Martínez-Montes et al., “Concurrent EEG/fMRI analysis by multiway Partial Least Squares,” Neuroimage 22(3), 1023–1034 (2004) [doi:10.1016/j.neuroimage.2004.03.038].

29. E. Acar et al., “Multiway analysis of epilepsy tensors,” Bioinformatics 23(13), i10–i18 (2007) [doi:10.1093/bioinformatics/btm210].

30. V. Calhoun, “Data-driven approaches for identifying links between brain structure and function in health and disease,” Dialogues Clin Neurosci 20(2), 87–99 (2018).

31. G. H. Klem et al., “The ten-twenty electrode system of the International Federation,” Electroencephalogr Clin Neurophysiol 52(3), 3–6 (1999).

32. W. D. Gaillard et al., “Developmental aspects of language processing: fMRI of verbal fluency in children and adults,” Human Brain Mapping 18(3), 176–185 (2003) [doi:10.1002/hbm.10091].

33. N. Paquette et al., “Developmental patterns of expressive language hemispheric lateralization in children, adolescents and adults using functional near-infrared spectroscopy,” Neuropsychologia 68, 117–125 (2015) [doi:10.1016/j.neuropsychologia.2015.01.007].

34. A. Gallagher, J. Tremblay, and P. Vannasing, “Language mapping in children using resting-state functional connectivity: comparison with a task-based approach,” Journal of Biomedical Optics 21(12), 125006 (2016) [doi:10.1117/1.JBO.21.12.125006].

35. J. Tremblay et al., “LIONirs: flexible Matlab toolbox for fNIRS data analysis,” Journal of Neuroscience Methods 370, 109487, Elsevier (2022) [doi:10.1016/j.jneumeth.2022.109487].

36. A. Villringer and U. Dirnagl, “Coupling of brain activity and cerebral blood flow: basis of functional neuroimaging,” Cerebrovasc Brain Metab Rev 7(3), 240–276 (1995).

37. A. Aarabi and T. J. Huppert, “Characterization of the relative contributions from systemic physiological noise to whole-brain resting-state functional near-infrared spectroscopy data using single-channel independent component analysis,” Neurophotonics 3(2), 025004 (2016) [doi:10.1117/1.NPh.3.2.025004].

38. G. H. Glover, “Deconvolution of Impulse Response in Event-Related BOLD fMRI1,” Neuroimage 9(4), 416–429 (1999) [doi:10.1006/nimg.1998.0419].

39. L. Gagnon et al., “Improved recovery of the hemodynamic response in diffuse optical imaging using short optode separations and state-space modeling,” Neuroimage 56(3), 1362–1371, Elsevier Inc. (2011) [doi:10.1016/j.neuroimage.2011.03.001].

40. L. Gagnon et al., “Short separation channel location impacts the performance of short channel regression in NIRS,” Neuroimage 59(3), 2518–2528 (2012) [doi:10.1016/j.neuroimage.2011.08.095].

41. M. Plank, “Ocular correction ICA,” Brain Products Press Release 49 (2013).

42. M. Mørup et al., “Parallel Factor Analysis as an exploratory tool for wavelet transformed event-related EEG,” Neuroimage 29(3), 938–947 (2006) [doi:10.1016/j.neuroimage.2005.08.005].

43. C. A. Andersson and R. Bro, “The N-way Toolbox for MATLAB,” Chemometrics and Intelligent Laboratory Systems 52(1), 1–4 (2000) [doi:10.1016/S0169-7439(00)00071-X].

44. K. T. Sweeney et al., “A Methodology for Validating Artifact Removal Techniques for Physiological Signals,” IEEE Transactions on Information Technology in Biomedicine 16(5), 918–926 (2012) [doi:10.1109/TITB.2012.2207400].

45. F. Scholkmann et al., “How to detect and reduce movement artifacts in near-infrared imaging using moving standard deviation and spline interpolation,” Physiological Measurement 31(5), 649–662 (2010) [doi:10.1088/0967-3334/31/5/004].

46. S. Liu et al., “Empirical mode decomposition applied to tissue artifact removal from respiratory signal,” in 2008 30th Annual International Conference of the IEEE Engineering in Medicine and Biology Society, pp. 3624–3627, IEEE (2008) [doi:10.1109/IEMBS.2008.4649991].

47. L. Kocsis, P. Herman, and A. Eke, “The modified Beer–Lambert law revisited,” Physics in Medicine and Biology 51(5), N91–N98 (2006) [doi:10.1088/0031-9155/51/5/N02].

48. F. Scholkmann and M. Wolf, “General equation for the differential pathlength factor of the frontal human head depending on wavelength and age,” Journal of Biomedical Optics 18(10), 105004, SPIE-Intl Soc Optical Eng (2013) [doi:10.1117/1.jbo.18.10.105004].

49. K. J. Friston et al., Statistical Parametric Mapping: The Analysis of Functional Brain Images, Academic Press, London, UK (2011).

50. M. L. Schroeter et al., “Towards a standard analysis for functional near-infrared imaging,” Neuroimage 21(1), 283–290 (2004) [doi:10.1016/j.neuroimage.2003.09.054].

51. Y. Minagawa-Kawai et al., “Optical Brain Imaging Reveals General Auditory and Language-Specific Processing in Early Infant Development,” Cerebral Cortex 21(2), 254–261 (2011) [doi:10.1093/cercor/bhq082].

52. J. Cohen, Statistical Power Analysis for the Behavioral Sciences, Second, Lawrence Erlbaum Associates, Inc., New York, NY (1988).

53. T. T. Liu, “Neurovascular factors in resting-state functional MRI,” Neuroimage 80, 339–348 (2013) [doi:10.1016/J.NEUROIMAGE.2013.04.071].

54. M. Mørup, “Applications of tensor (multiway array) factorizations and decompositions in data mining,” Wiley Interdisciplinary Reviews: Data Mining and Knowledge Discovery 1(1), 24–40 (2011) [doi:10.1002/widm.1].

55. S. Tak et al., “Dynamic causal modelling for functional near-infrared spectroscopy,” Neuroimage 111, 338–349 (2015) [doi:10.1016/j.neuroimage.2015.02.035].

56. M. A. Yücel et al., “Best practices for fNIRS publications,” Neurophotonics 8(01), 012101, International Society for Optics and Photonics (2021) [doi:10.1117/1.nph.8.1.012101].

57. E. Martínez-Montes, J. M. Sánchez-Bornot, and P. A. Valdés-Sosa, “Penalized PARAFAC analysis of spontaneous EEG recordings,” Stat Sin 18(4), 1449–1464 (2008).

58. E. Martínez-Montes et al., “Identifying Complex Brain Networks Using Penalized Regression Methods,” Journal of Biological Physics 34(3–4), 315–323 (2008) [doi:10.1007/s10867-008-9077-0].

59. D. Poeppel et al., “Towards a New Neurobiology of Language,” Journal of Neuroscience 32(41), 14125–14131 (2012) [doi:10.1523/JNEUROSCI.3244-12.2012].

60. S. Makeig, “Dynamic Brain Sources of Visual Evoked Responses,” Science (1979) 295(5555), 690–694 (2002) [doi:10.1126/science.1066168].

61. N. D. Sidiropoulos and R. Bro, “On the uniqueness of multilinear decomposition of N-way arrays,” Journal of Chemometrics 14(3), 229–239 (2000) [doi:10.1002/1099-128X(200005/06)14:3<229::AID-CEM587>3.0.CO;2-N].

62. W. Deburchgraeve et al., “Neonatal seizure localization using PARAFAC decomposition,” Clinical Neurophysiology 120(10), 1787–1796 (2009) [doi:10.1016/j.clinph.2009.07.044].

63. T. J. Huppert, “Commentary on the statistical properties of noise and its implication on general linear models in functional near-infrared spectroscopy,” Neurophotonics 3(1), 010401 (2016) [doi:10.1117/1.NPh.3.1.010401].

64. M. Hu and H. Liang, “A copula approach to assessing Granger causality,” Neuroimage 100, 125–134 (2014) [doi:10.1016/j.neuroimage.2014.06.013].

65. W. A. Freiwald et al., “Testing non-linearity and directedness of interactions between neural groups in the macaque inferotemporal cortex,” Journal of Neuroscience Methods 94(1), 105–119 (1999) [doi:10.1016/S0165-0270(99)00129-6].

66. N. M. Faber, R. Bro, and P. K. Hopke, “Recent developments in CANDECOMP/PARAFAC algorithms: A critical review,” in Chemometrics and Intelligent Laboratory Systems 65(1), pp. 119–137 (2003) [doi:10.1016/S0169-7439(02)00089-8].

67. G. Jasdzewski et al., “Differences in the hemodynamic response to event-related motor and visual paradigms as measured by near-infrared spectroscopy,” Neuroimage 20(1), 479–488, Academic Press Inc. (2003) [doi:10.1016/S1053-8119(03)00311-2].

68. T. Sato et al., “Reduction of global interference of scalp-hemodynamics in functional near-infrared spectroscopy using short distance probes,” Neuroimage 141, 120–132 (2016) [doi:10.1016/j.neuroimage.2016.06.054].

69. J. Tremblay et al., “Comparison of source localization techniques in diffuse optical tomography for fNIRS application using a realistic head model,” Biomedical Optics Express 9(7), 448–454 (2018) [doi:10.1364/BOE.9.002994].

70. M. Ontivero-Ortega, Y. Garcia-Puente, and E. Martínez-Montes, “Comparison of classifiers to detect epileptic seizures via PARAFAC decomposition,” in IFMBE Proceedings 49, pp. 500–503, Springer, Cham (2015) [doi:10.1007/978-3-319-13117-7_128].

71. K. Nazarpour et al., “Parallel space-time-frequency decomposition of EEG signals for brain computer interfacing; Parallel space-time-frequency decomposition of EEG signals for brain computer interfacing” (2006).

72. A. Cichocki et al., “Noninvasive BCIs: Multiway Signal-Processing Array Decompositions,” Computer (Long Beach Calif) 41(10), 34–42 (2008) [doi:10.1109/MC.2008.431].

73. A. Eliseyev et al., “L1-Penalized N-way PLS for subset of electrodes selection in BCI experiments,” Journal of Neural Engineering 9(4) (2012) [doi:10.1088/1741-2560/9/4/045010].

74. A. Eliseyev and T. Aksenova, “Recursive N-Way Partial Least Squares for Brain-Computer Interface,” PLoS ONE 8(7), 69962 (2013) [doi:10.1371/journal.pone.0069962].

